# Meta-analysis of problematic alcohol use in 435,563 individuals identifies 29 risk variants and yields insights into biology, pleiotropy and causality

**DOI:** 10.1101/738088

**Authors:** Hang Zhou, Julia M. Sealock, Sandra Sanchez-Roige, Toni-Kim Clarke, Daniel Levey, Zhongshan Cheng, Boyang Li, Renato Polimanti, Rachel L. Kember, Rachel Vickers Smith, Johan H. Thygesen, Marsha Y. Morgan, Stephen R. Atkinson, Mark R. Thursz, Mette Nyegaard, Manuel Mattheisen, Anders D. Børglum, Emma C. Johnson, the VA Million Veteran Program, Amy C. Justice, Abraham A. Palmer, Andrew McQuillin, Lea K. Davis, Howard J. Edenberg, Arpana Agrawal, Henry R. Kranzler, Joel Gelernter

**Affiliations:** Department of Psychiatry, Yale School of Medicine, New Haven, CT, USA; Veterans Affairs Connecticut Healthcare System, West Haven, CT, USA; Vanderbilt Genetics Institute, Vanderbilt University Medical Center, Nashville, TN, USA; Division of Medical Genetics, Department of Medicine, Vanderbilt University Medical Center, Nashville, TN, USA; Department of Psychiatry, University of California San Diego, La Jolla, CA, USA; Division of Psychiatry, University of Edinburgh, Edinburgh, UK; Department of Biostatistics, Yale School of Public Health, New Haven, CT, USA; Department of Genetics, University of Pennsylvania Perelman School of Medicine, Philadelphia, PA, USA; Crescenz Veterans Affairs Medical Center, Philadelphia, PA, USA; University of Louisville School of Nursing, Louisville, KY, USA; Division of Psychiatry, University College London, London, UK; UCL Institute for Liver & Digestive Health, Division of Medicine, Royal Free Campus, University College London, London, UK; Department of Metabolism, Digestion & Reproduction, Imperial College London, London, UK; Department of Biomedicine, Aarhus University, Aarhus, Denmark; Centre for Integrative Sequencing, iSEQ, Aarhus University, Aarhus, Denmark; The Lundbeck Foundation Initiative for Integrative Psychiatric Research, iPSYCH, Denmark; Center for Genomics and Personalized Medicine, Aarhus, Denmark; Department of Psychiatry, Psychosomatics and Psychotherapy, University of Würzburg, Würzburg, Germany; Department of Clinical Neuroscience, Karolinska Institutet, Stockholm, Sweden; Department of Psychiatry, Washington University School of Medicine, St. Louis, MO, USA; Department of Internal Medicine, Yale School of Medicine, New Haven, CT, USA; Center for Interdisciplinary Research on AIDS, Yale School of Public Health, New Haven, CT, USA; Institute for Genomic Medicine, University of California San Diego, La Jolla, CA, USA; Department of Psychiatry and Behavioral Sciences, Vanderbilt University Medical Center, Nashville, TN, USA; Department of Biochemistry and Molecular Biology, Indiana University School of Medicine, Indianapolis, IN, USA; Department of Medical and Molecular Genetics, Indiana University School of Medicine, Indianapolis, IN, USA; Department of Psychiatry, University of Pennsylvania Perelman School of Medicine, Philadelphia, PA, USA; Departments of Genetics and Neuroscience, Yale University School of Medicine

## Abstract

Problematic alcohol use (PAU) is a leading cause of death and disability worldwide. Although genome-wide association studies (GWASs) have identified PAU risk genes, the genetic architecture of this trait is not fully understood. We conducted a proxy-phenotype meta-analysis of PAU combining alcohol use disorder and problematic drinking in 435,563 European-ancestry individuals. We identified 29 independent risk variants, 19 of them novel. PAU was genetically correlated with 138 phenotypes, including substance use and psychiatric traits. Phenome-wide polygenic risk score analysis in an independent biobank sample (BioVU, *n*=67,589) confirmed the genetic correlations between PAU and substance use and psychiatric disorders. Genetic heritability of PAU was enriched in brain and in genomic conserved and regulatory regions. Mendelian randomization suggested causal effects on liability to PAU of substance use, psychiatric status, risk-taking behavior, and cognitive performance. In summary, this large PAU meta-analysis identified novel risk loci and revealed genetic relationships with numerous other outcomes.

## Introduction

Alcohol use and alcohol use disorder (AUD) are leading causes of death and disability worldwide [1]. Genome-wide association studies (GWAS) of AUD and problematic drinking measured by different assessments have identified potential risk genes primarily in European populations [2–5]. Quantity-frequency measures of drinking, for example the Alcohol Use Disorders Identification Test–Consumption (AUDIT-C), which sometimes reflect alcohol consumption in the normal range, differ genetically from AUD and measures of problematic drinking (e.g., the Alcohol Use Disorders Identification Test–Problems [AUDIT-P]), and show a divergent set of genetic correlations [3, 4]. The estimated SNP-based heritability (*h*^2^) of AUD ranges from 5.6% to 10.0% [2–5]. To date, more than ten risk variants have been significantly associated with AUD and AUDIT-P (p < 5 × 10^−8^). Variants mapped to several risk genes have been detected in multiple studies, including *ADH1B* (Alcohol Dehydrogenase 1B), *ADH1C* (Alcohol Dehydrogenase 1C), *ALDH2* (Aldehyde Dehydrogenase 2, only in some Asian samples), *SLC39A8* (Solute Carrier Family 39 Member 8), *GCKR* (Glucokinase Regulator), and *CRHR1* (Corticotropin Releasing Hormone Receptor 1). In the context of the known extensive polygenicity underlying AUD and AUDIT-P, we anticipate that additional significant risk loci can be identified by increasing sample size; this is the pattern for GWAS of heterogenous complex traits in general also. We can characterize both AUD itself and AUDIT-P, as “problematic alcohol use” (PAU). To identify additional risk variants and enhance our understanding of the genetic architecture of PAU, we conducted genome-wide meta-analysis of AUD and AUDIT-P in 435,563 individuals of European ancestry. The understanding of the genetic architecture of PAU in African populations is far behind than Europeans; the largest sample published so far is 56,648 in MVP [3] and results have not moved beyond a single genomic region that includes *ADH1B*. This study only focused on European samples because we cannot achieve a substantial increment in African-ancestry subjects over previous studies.

## Results

Figure 1 provides an overview of the meta-analysis of the 4 major datasets. The first is the GWAS of AUD in European Americans (EA) from the Million Veteran Program (MVP) [6] (herein designated “MVP phase1”), comprised 202,004 individuals phenotyped for AUD (*n*_case_ = 34,658, *n*_control_ = 167,346, *n*_effective_ = 114,847) using International Classification of Diseases (ICD) codes [3]. The second, MVP Phase2, included an additional 65,387 EA individuals from MVP (*n*_case_ = 11,337, *n*_control_ = 54,050, *n*_effective_ = 37,485) not previously analyzed. The third dataset is a GWAS of DSM-IV alcohol dependence (AD) from the Psychiatric Genomics Consortium (PGC), which included 46,568 European participants (*n*_case_ = 11,569, *n*_control_ = 34,999, *n*_effective_ = 26,853) [2]. The fourth dataset is a GWAS of Alcohol Use Disorders Identification Test–Problems (AUDIT-P; a measure of problematic drinking) scores from a UK Biobank sample (UKB) [7] that included 121,604 European participants [4].

**Figure 1.**
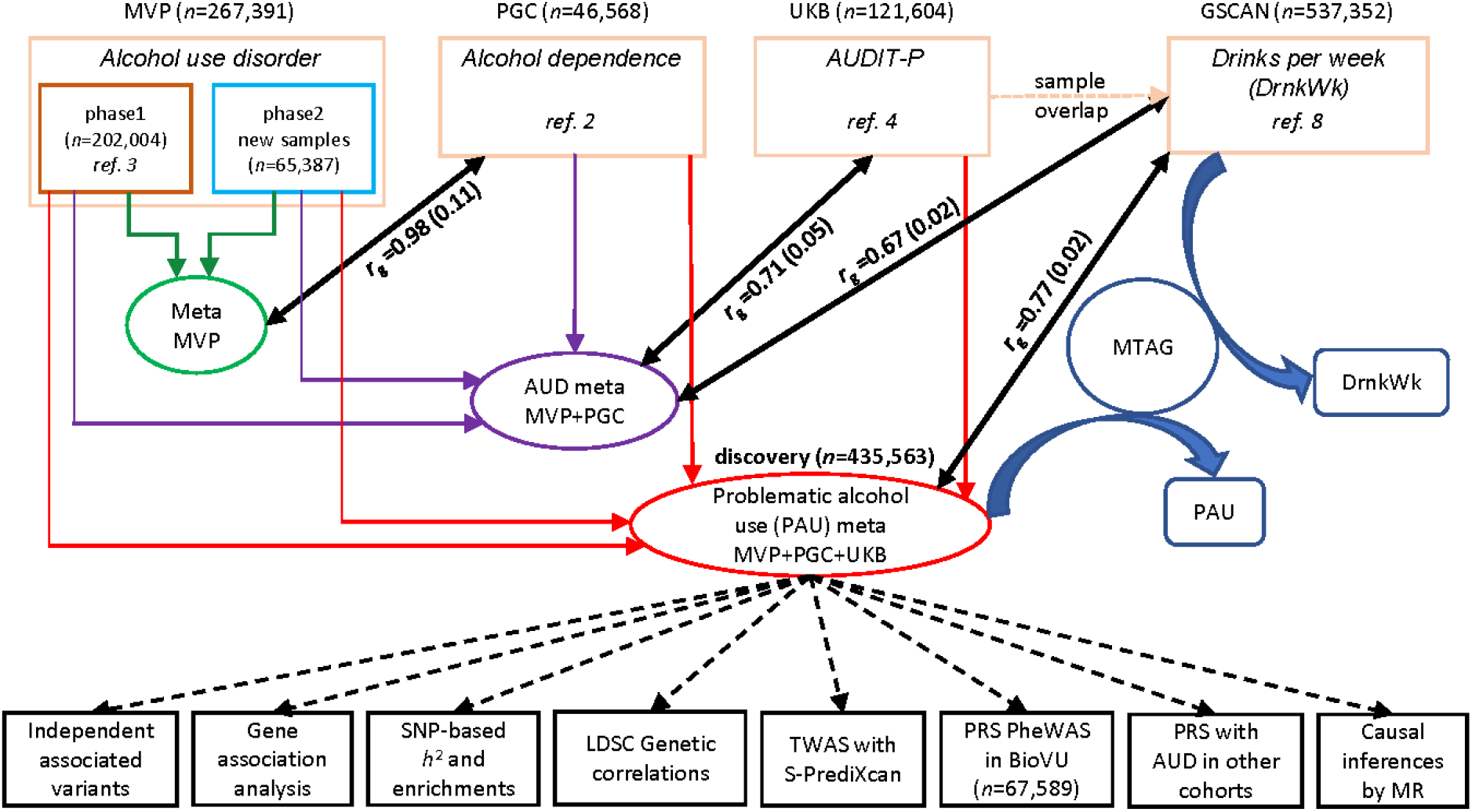
Overview of the analysis. Four datasets were meta-analyzed as the discovery sample for problematic alcohol use (PAU) including MVP phase1, MVP phase2, PGC, and UK Biobank (UKB). MVP phase1 and phase2 were meta-analyzed, and the result was used for testing the genetic correlation with PGC alcohol dependence. An intermediary meta-analysis (AUD meta) combining MVP phase1, phase2, and PGC was then conducted to measure the genetic correlation with UKB AUDIT-P. Due to the sample overlap between UKB and GSCAN, we used the AUD meta-analysis for Mendelian Randomization (MR) analysis rather than the PAU (i.e., the second) meta-analysis. MTAG, which used the summary data from PAU and DrnkWk (drinks per week) in GSCAN (without 23andMe samples as those data were not made available) as input to increase the power for each trait without introducing bias from sample overlap, returned summary results for PAU and DrnkWk separately.

The genetic correlation (*r*_g_) between MVP phase1 AUD and PGC AD was 0.965 (se = 0.15, p = 1.21 × 10^−10^) [3]. *R*_*g*_ between the entire MVP (meta-analysis of phase1 and phase2) and PGC increased to 0.98 (se = 0.11, p = 1.99 × 10^−19^), justifying the meta-analysis of AUD across the three datasets (*n*_effective_ = 179,185). 24 risk variants in 23 loci were detected in this intermediary meta-analysis (Supplementary Figure 1, Supplementary Table 1). The *r*_g_ between UKB AUDIT-P and AUD (MVP+PGC) was 0.71 (se = 0.05, p = 8.15 × 10^−52^), and the polygenic risk score (PRS) of AUD was associated with AUDIT-P in UKB (best p-value threshold PT_best_ = 0.001, R^2^ = 0.25%, p = 3.28 × 10^−41^, Supplementary Table 2, Supplementary Figure 2), justifying the proxy-phenotype meta-analysis of problematic alcohol use (PAU) across all four datasets. The total sample size was 435,563 in the discovery analysis (*n*_effective_ = 300,789).

### Association results for PAU

Of 42 lead variants (mapping to 27 loci, Supplementary Figure 3, and Supplementary Table 3) that were genome-wide significant (GWS) for PAU, 29 were independently associated after conditioning on lead SNPs in the regions (see below and Table 1). Ten variants were previously identified through the same index SNPs or tagged SNPs, located in or near the following genes: *GCKR*, *SIX3*, *KLB*, *ADH1B*, *ADH1C*, *SLC39A8*, *DRD2*, and *FTO* [2–5]. Thus, 19 variants reported here were novel, of which 11 were located in gene regions, including *PDE4B* (Phosphodiesterase 4B), *THSD7B* (Thrombospondin Type 1 Domain Containing 7B), *CADM2* (Cell Adhesion Molecule 2), *ADH1B* (different from the locus identified previously), *DPP6* (Dipeptidyl Peptidase Like 6), *SLC39A13* (Solute Carrier Family 39 Member 13), *TMX2* (Thioredoxin Related Transmembrane Protein 2), *ARID4A* (AT-Rich Interaction Domain 4A), *C14orf2* (Chromosome 14 open reading frame 2), *TNRC6A* (Trinucleotide Repeat Containing Adaptor 6A), and *FUT2* (Fucosyltransferase 2). A novel rare *ADH1B* variant, rs75967634 (p = 1.07 × 10^−9^, with a minor allele frequency of 0.003), which causes a substitution of histidine for arginine, is in the same codon as rs2066702 (a well-known variant associated with AUD in African populations[3, 8], but not polymorphic in European populations).This latter association is independent from rs1229984 in *ADH1B* and rs13125415 (a tag SNP of rs1612735 in MVP phase1 [3]) in *ADH1C*. The identification of rs75967634 demonstrates the present study’s greater power to detect risk variants in this region, beyond the frequently reported *ADH1B\**rs1229984.

**Table 1.** Genome-wide significant associations for PAU.

| Chr | Pos (hg19) | rsID | Gene | A1 | A2 | EAf | Z | P | Direction |
| --- | --- | --- | --- | --- | --- | --- | --- | --- | --- |
| 1 | 66419905 | **rs6421482** | *PDE4B*<sup>a</sup> | A | G | 0.4363 | -6.315 | 2.7×10<sup>-10</sup> | ---- |
| 1 | 73848610 | **rs61767420** | [] | A | G | 0.3999 | 5.714 | 1.11×10<sup>-8</sup> | ++++ |
| 2 | 27730940 | rs1260326 | *GCKR*<sup>a</sup> | T | C | 0.4033 | -9.296 | 1.45×10<sup>-20</sup> | --+- |
| 2 | 45141180 | rs494904 | *SIX3*<sup>b</sup> | T | C | 0.5961 | -7.926 | 2.26×10<sup>-15</sup> | ---- |
| 2 | 58042241 | **rs1402398** | *VRK2*<sup>b</sup> | A | G | 0.6274 | 7.098 | 1.27×10<sup>-12</sup> | ++++ |
| 2 | 104134432 | **rs9679319** | [] | T | G | 0.4797 | -6.01 | 1.86×10<sup>-9</sup> | ---- |
| 2 | 138264231 | **rs13382553** | *THSD7B*<sup>a</sup> | A | G | 0.766 | -6.001 | 1.97×10<sup>-9</sup> | ---- |
| 2 | 227164653 | **rs2673136** | *IRS1*<sup>b</sup> | A | G | 0.6387 | -5.872 | 4.31×10<sup>-9</sup> | ---- |
| 3 | 85513793 | **rs62250713** | *CADM2*<sup>a</sup> | A | G | 0.368 | 6.049 | 1.46×10<sup>-9</sup> | ++++ |
| 4 | 39404872 | rs13129401 | *KLB*<sup>b</sup> | A | G | 0.4532 | -8.906 | 5.29×10<sup>-19</sup> | ---- |
| 4 | 100229016 | **rs75967634** | *ADH1B*<sup>a</sup> | T | C | 0.003 | -6.098 | 1.07×10<sup>-9</sup> | --?- |
| 4 | 100239319 | rs1229984 | *ADH1B*<sup>a</sup> | T | C | 0.0302 | -22 | 2.9×10<sup>-107</sup> | ---? |
| 4 | 100270452 | rs13125415 | *ADH1C*<sup>a</sup> | A | G | 0.5849 | -9.073 | 1.16×10<sup>-19</sup> | ---- |
| 4 | 103198082 | rs13135092 | *SLC39A8*<sup>a</sup> | A | G | 0.9192 | 11.673 | 1.75×10<sup>-31</sup> | ++++ |
| 7 | 153489074 | **rs2533200** | *DPP6*<sup>a</sup> | C | G | 0.5163 | -5.631 | 1.79×10<sup>-8</sup> | ---- |
| 8 | 57424874 | **rs2582405** | *PENK*<sup>b</sup> | T | C | 0.237 | 5.751 | 8.86×10<sup>-9</sup> | ++++ |
| 10 | 72907951 | **rs7900002** | *UNC5B*<sup>b</sup> | T | G | 0.6012 | -5.503 | 3.74×10<sup>-8</sup> | --+- |
| 10 | 110537834 | rs56722963 | [] | T | C | 0.2551 | -6.374 | 1.85×10<sup>-10</sup> | ---- |
| 11 | 47423920 | **rs10717830** | *SLC39A13*<sup>a</sup> | G | GT | 0.674 | 6.422 | 1.34×10<sup>-10</sup> | ++++ |
| 11 | 57480623 | **rs576859** | *TMX2*<sup>a</sup> | A | C | 0.3272 | 5.67 | 1.43×10<sup>-8</sup> | +++? |
| 11 | 113357710 | rs138084129 | *DRD2*<sup>b</sup> | A | AATAT | 0.6274 | 7.824 | 5.13×10<sup>-15</sup> | ++++ |
| 11 | 113443753 | rs6589386 | *DRD2*<sup>b</sup> | T | C | 0.4323 | -7.511 | 5.88×10<sup>-14</sup> | ---- |
| 11 | 121542923 | **rs1783835** | *SORL1*<sup>b</sup> | A | G | 0.4569 | -5.979 | 2.24×10<sup>-9</sup> | ---- |
| 12 | 51903860 | **rs12296477** | *SLC4A8*<sup>b</sup> | C | G | 0.5469 | 5.484 | 4.15×10<sup>-8</sup> | ++++ |
| 14 | 58765903 | **rs61974485** | *ARID4A*<sup>a</sup> | T | C | 0.2646 | 5.506 | 3.67×10<sup>-8</sup> | ++++ |
| 14 | 104355883 | **rs8008020** | *C14orf2*<sup>a</sup> | T | C | 0.4175 | 6.062 | 1.35×10<sup>-9</sup> | ++++ |
| 16 | 24693048 | **rs72768626** | *TNRC6A*<sup>a</sup> | A | G | 0.9448 | 5.591 | 2.26×10<sup>-8</sup> | ++++ |
| 16 | 53820813 | rs9937709 | *FTO*<sup>a</sup> | A | G | 0.585 | 6.602 | 4.06×10<sup>-11</sup> | ++++ |
| 19 | 49206417 | **rs492602** | *FUT2*<sup>a</sup> | A | G | 0.5076 | -6.143 | 8.08×10<sup>-10</sup> | ---- |

Listed are the 29 independent variants that were genome-wide significant. Variants labeled in bold are novel associations with PAU. A1, effect allele; A2, other allele; EAF, effective allele frequency; Directions are for the A1 allele in MVP phase1, MVP phase2, PGC, and UKB datasets.
<sup>a</sup>Protein-coding gene contains the lead SNP,
<sup>b</sup>Protein-coding gene nearest to the lead SNP.

Moderate genetic correlation between AUD and alcohol consumption, and also pervasive pleiotropic effects of SNPs, were demonstrated previously [2–4]. Some of the novel variants (10 out of 19) identified in this study were also associated with other alcohol-related traits, including AUDIT-C score [3], total AUDIT score [4], and drinks per week (DrnkWk) from the GSCAN study [9] (described below and Supplementary Table 3). Rs1402398, close to *VRK2*, was associated with AUDIT-C score (tagged by rs2683616) [3]; rs492602 in *FUT2* was associated with DrnkWk [9] and total AUDIT score [4]; and rs6421482, rs62250713, rs2533200, rs10717830, rs1783835, rs12296477, rs61974485, and rs72768626 were associated with DrnkWk directly or through tag SNPs in high linkage disequilibrium (LD) [9]. Analysis conditioned on DrnkWk shows that 11 of the 29 independent variants were independently associated with PAU (i.e., not mediated by DrnkWk) (Supplementary Table 3).

#### Gene-based association analysis

identified 66 genes that were associated with PAU at GWS (p < 2.64 × 10^−6^, Supplementary Table 4). *DRD2*, which has been extensively studied in many fields of neuroscience, was among these 66 genes and had been reported in both UKB [4] and MVP phase1 [3]. Among the 66 genes, 46 are novel, including *ADH4* (Alcohol Dehydrogenase 4), *ADH5* (Alcohol Dehydrogenase 5), and *ADH7* (Alcohol Dehydrogenase 7), extending alcohol metabolizing gene associations beyond the well-known *ADH1B* and *ADH1C*; *SYNGAP1* (Synaptic Ras GTPase Activating Protein 1), *BDNF* (Brain-Derived Neurotrophic Factor), and others. Certain genes show associations with multiple traits including previous associations with AUDIT-C (4 genes in MVP phase1, 12 genes in UKB), total AUDIT score (19 genes in UKB), and DrnkWk (46 genes in GSCAN, which includes results for DrnkWk after MTAG [10] analysis).

Examination of the 66 associated genes for known drug-gene interactions through the Drug Gene Interaction Database v3.0.2 [11] showed 327 interactions between 16 genes and 325 drugs (Supplementary Table 5). Of these 16 genes with interactions, *DRD2* had the most drug interactions (*n* = 177), followed by *BDNF* (*n* = 68) and *PDE4B* (*n* = 36).

### SNP-based *h*^2^ and partitioning heritability enrichment

We used LD Score Regression (LDSC) [12] to estimate SNP-based *h*^2^ in the different datasests and the meta-analyses (Figure 2). Because of the unbalanced case/control ratio, we used effective sample size instead of actual sample size in MVP (following the PGC AD GWAS [2]). The *h*^2^ of PAU (the meta result) was 0.068 (se = 0.004). The *h*^2^ of AUD in the MVP meta-analysis (phases 1 and 2) was 0.095 (se = 0.006), and was 0.094 (se = 0.005) in the meta-analysis combining MVP and PGC.

**Figure 2.**
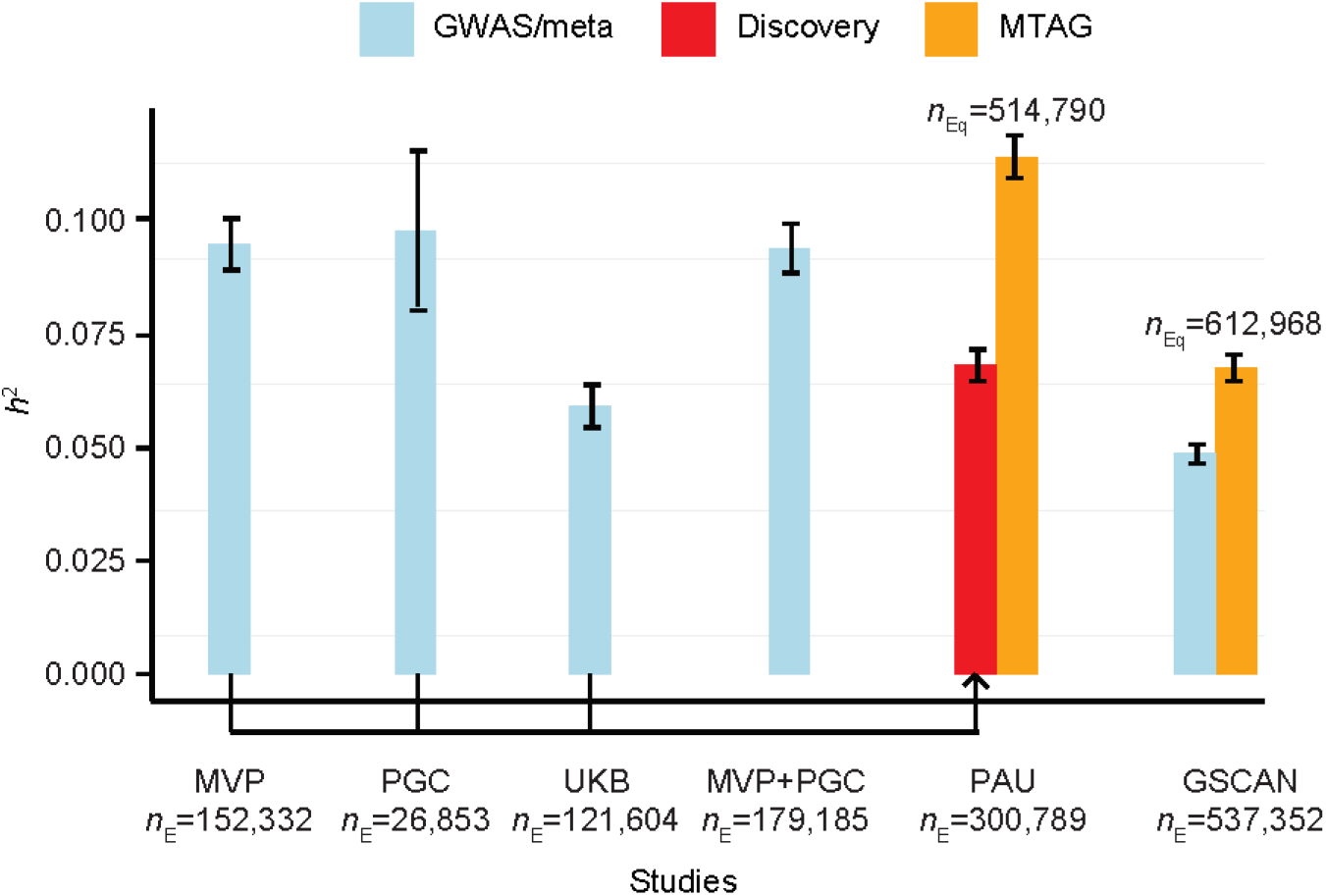
Estimated SNP-based *h*^2^. Blue bars show *h*^2^ results for single datasets or meta-analysis between datasets, from published studies or analyzed here. Red bar shows *h*^2^ for the PAU discovery meta-analysis. Orange bars show *h*^2^ results of MTAG analysis for PAU in the discovery sample and DrnkWk from GSCAN. Effective sample sizes (*n*_E_) were used in LDSC. *n*_Eq_ is the GWAS-equivalent sample size reported by MTAG.

Partitioning heritability enrichment analyses using LDSC [13, 14] showed the most significantly enriched cell type group to be central nervous system (CNS, p = 3.53 × 10^−9^), followed by adrenal and pancreas (p = 1.89 × 10^−3^), and immune and hematopoietic (p = 3.82 × 10^−3^, Supplementary Figure 4). Significant enrichments were also observed in six baseline annotations, including conserved regions, conserved regions with 500bp extended (ext), fetal DHS (DNase I hypersensitive sites) ext, weak enhancers ext, histone mark H3K4me1 ext, and TSS (transcription start site) ext (Supplementary Figure 5). We also investigated heritability enrichments using Roadmap data, which contains six annotations (DHS, H3K27ac, H3K4me3, H3K4me1, H3K9ac, and H3K36me3) in a subset of 88 primary cell types and tissues [14, 15]. Significant enrichments were observed for H3K4me1 and DHS in fetal brain, and H3K4me3 in fetal brain and in brain germinal matrix (Supplementary Table 6). Although no heritability enrichment was observed in tissues using gene expression data from GTEx [16], the top nominally enriched tissues were all in brain (Supplementary Figure 6).

### Functional enrichments

MAGMA tissue expression analysis [17, 18] using GTEx showed significant enrichments in several brain tissues including cerebellum and cortex (Supplementary Figure 7). Although no enrichment was observed via MAGMA gene-set analysis using gene-based p-values of all protein-coding genes, the 152 genes prioritized by positional, expression quantitative trait loci (eQTL), and chromatin interaction mapping were enriched in several gene sets, including ethanol metabolic processes (Supplementary Table 7).

### Genetic correlations with other traits

We estimated the genetic correlations between PAU and 715 publicly available sets of GWAS summary statistics which included 228 published sets and 487 unpublished sets from the UK Biobank. After Bonferroni correction (p < 6.99 × 10^−5^), 138 traits were significantly correlated with PAU (Supplementary Table 8). Among the 26 published traits, drinks per week showed the highest *r*_g_ with PAU (*r*_g_ = 0.77, se = 0.02, p = 3.25 × 10^−265^), consistent with the overall quantity of alcohol consumed being a key domain of PAU [5, 19]. Several smoking traits and lifetime cannabis use were positively genetically correlated with PAU, consistent with the high comorbidity between alcohol and other substance use disorders in the general population [20]. Among psychiatric disorders, major depressive disorder (MDD, *r*_g_ = 0.39, se = 0.03, p = 1.43 × 10^−40^) showed the highest genetic correlation with PAU, extending the evidence for the shared genetic contribution to MDD and alcohol-related traits [21, 22]. PAU was positively genetically correlated with risk-taking behavior, insomnia, lung cancer, and other traits, and negatively correlated with cognitive traits and parents’ age at death. These finding are in line with the known adverse medical, psychiatric, and social consequences of problem drinking (Figure 3).

**Figure 3.**
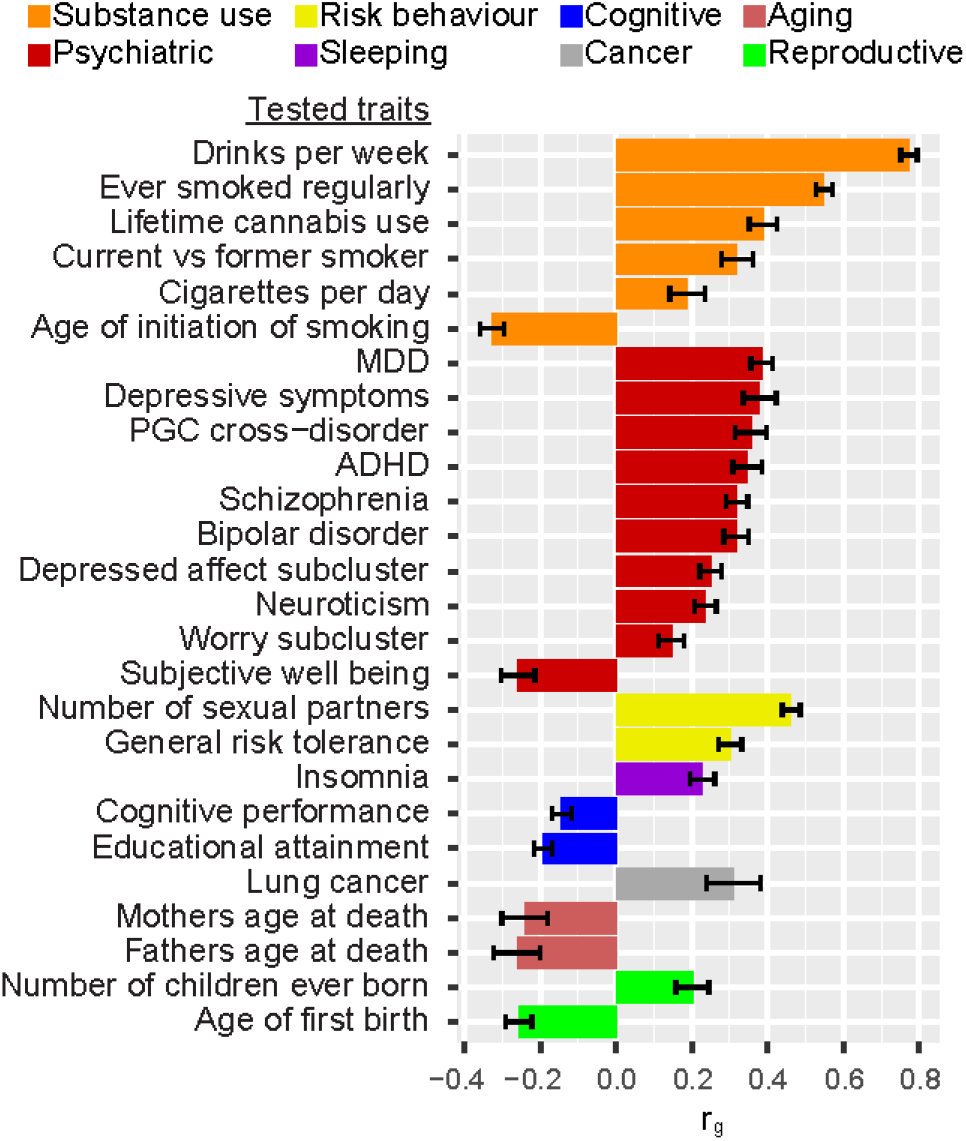
Genetic correlations with published traits. Of 228 published traits, 26 were genetically correlated with PAU after Bonferroni correction (p < 0.05/715). MDD, major depressive disorder; ADHD, attention deficit hyperactivity disorder.

### Transcriptomic analyses

We used S-PrediXcan [23] to predict gene expression and the mediating effects of variation on gene expression on PAU. Forty-eight tissues from GTEx [16] release v7 and whole blood samples from the Depression Genes and Networks study (DGN) [24] were analyzed as reference transcriptomes (Supplementary Table 9). After Bonferroni correction, 103 gene-tissue associations were significant, representing 39 different genes, some of which were identified in multiple tissues (Supplementary Table 10). For example, *C1QTNF4* (C1q and tumor necrosis factor related protein 4) was detected in 18 tissues, including brain, gastrointestinal, adipose, and liver. None of the four significant alcohol dehydrogenase genes (*ADH1A*, *ADH1B*, *ADH4*, and *ADH5*) was associated with expression in brain tissue, but they were associated with expression in other tissues --adipose, thyroid, gastrointestinal and heart. This might be due to the generally low expression level of these genes in brain [25]. These cross-tissue associations indicate that there are widespread functional consequences of PAU-risk-associated genetic variation at the expression level.

Although the sample size for tissues used for eQTL analysis limits our ability to detect associations, there are substantial common eQTLs across tissues [16]. Integrating evidence from multiple tissues can increase power to detect genes relative to the tissues tested individually, at least for shared eQTLs. We applied S-MultiXcan [26] to the summary data for PAU using all 48 GTEx tissues as reference transcriptomic data. The expression of 34 genes was significantly associated with PAU, including *ADH1B*, *ADH4*, *ADH5*, *C1QTNF4*, *GCKR*, and *DRD2* (Supplementary Table 11). Among the 34 genes, 27 overlapped with genes detected by S-PrediXcan.

### PAU PRS for phenome-wide associations

We calculated PRS for PAU in 67,589 individuals of European descent from the Vanderbilt University Medical Center’s biobank, BioVU. We conducted a phenome-wide association study (PheWAS) of PRS for PAU adjusting for sex, age (calculated as the median age across an individual’s medical record), and top 10 principal components of ancestry. We standardized the PRS so that the odds ratios correspond to a standard deviation increase in the PRS. After Bonferroni correction, 31 of the 1,372 phenotypes tested were significantly associated with PAU PRS, including alcohol-related disorders (OR = 1.46, se = 0.03, p = 3.34 × 10^−40^), alcoholism (OR = 1.33, se = 0.03, p = 3.85 × 10^−28^), tobacco use disorder (OR = 1.21, se = 0.01, p = 2.71 × 10^−38^), 6 respiratory conditions, and 17 additional psychiatric conditions (Supplementary Figure 8, Supplementary Table 12).

### PAU PRS with AD in independent samples

We tested the association between PAU PRS and alcohol dependence in three independent samples: the iPSYCH group (*n*_case_ = 944, *n*_control_ = 11,408, *n*_effective_ = 3,487); University College London (UCL) Psych Array (*n*_case_ = 1,698, *n*_control_ = 1,228, *n*_effective_ = 2,851); and UCL Core Exome Array (*n*_case_ = 637, *n*_control_ = 9,189, *n*_effective_ = 2,383). The PAU PRSs were significantly associated with AD in all three samples, with the most variance explained in the UCL Psych Array sample, which includes the most cases (PT_best_ = 0.001, R^2^ = 2.12%, p = 8.64 × 10^−14^). In the iPSYCH group and UCL Core Exome Array samples, the maximal variance explained was 1.61% (PT_best_ = 0.3, p = 1.87 × 10^−22^), and 0.77% (PT_best_ = 5 × 10^−8^, p = 1.65 × 10^−7^), respectively (Supplementary Table 13).

### Mendelian Randomization

We tested the causal effects of liability to exposures on liability to AUD (MVP+PGC), rather than PAU: the UKB AUDIT-P GWAS was excluded to minimize sample overlap with other GWAS for putative exposures. We limited the exposures to those genetically correlated with PAU, and have more than 30 available instruments. There were only 24 independent variants for AUD; therefore the causal effects of liability to AUD on other traits (i.e., bidirectional) were not tested. Among the 13 tested exposures, 12 showed evidence of a causal effect on liability to AUD, the exception being cigarettes per day (Table 2). DrnkWk and ever smoked regularly have a positive causal effect on AUD risk by all 3 methods, without violating MR assumptions through horizontal pleiotropy (MR-Egger intercept p > 0.05). General risk tolerance was shown to be causally related to AUD risk, though the estimate could be biased due to horizontal pleiotropy (intercept p = 9.62 × 10^−3^). MDD, depressed affect neuroticism subcluster, worry neuroticism subcluster, number of sexual partners, and insomnia show evidence of positive causal effects on liability to AUD from at least one method, while cognitive performance and educational attainment show evidence of negative causal effects.

**Table 2.** Causal effects on liability to AUD (MVP+PGC) by MR.

| Exposure (#instruments) | Ref | IVW [27] |  | Weighted median [28] |  | MR-Egger [29] |  | MR-Egger intercept p |
| --- | --- | --- | --- | --- | --- | --- | --- | --- |
| | | $\beta$ (se) | p | $\beta$ (se) | p | $\beta$ (se) | p | |
| **DrnkWk (58)** | [9] | 0.89 (0.06) | **$1.80 \times 10^{-46}$** | 0.89 (0.08) | **$2.89 \times 10^{-26}$** | 0.91 (0.20) | **$3.80 \times 10^{-6}$** | 0.898 |
| **Ever smoked regularly (199)** | [9] | 0.32 (0.02) | **$8.72 \times 10^{-51}$** | 0.33 (0.02) | **$4.20 \times 10^{-43}$** | 0.26 (0.08) | **$1.21 \times 10^{-3}$** | 0.471 |
| Cigarettes per day (33) | [9] | 0.04 (0.06) | 0.475 | -0.10 (0.04) | 0.010 | -0.18 (0.09) | 0.034 | $1.27 \times 10^{-3}$ |
| **MDD (78)** | [30] | 0.14 (0.03) | **$8.42 \times 10^{-6}$** | 0.14 (0.03) | **$2.79 \times 10^{-6}$** | -0.17 (0.20) | 0.390 | 0.113 |
| Schizophrenia (110) | [31] | 0.04 (0.01) | **$2.47 \times 10^{-6}$** | 0.04 (0.01) | **$4.96 \times 10^{-6}$** | -0.05 (0.04) | 0.202 | 0.016 |
| **Depressed affect subcluster (56)** | [32] | 0.19 (0.06) | $1.75 \times 10^{-3}$ | 0.24 (0.05) | **$5.44 \times 10^{-6}$** | -0.20 (0.28) | 0.462 | 0.147 |
| Neuroticism (131) | [32] | 0.20 (0.04) | **$1.10 \times 10^{-7}$** | 0.20 (0.04) | **$1.10 \times 10^{-7}$** | -0.26 (0.16) | 0.097 | $2.64 \times 10^{-3}$ |
| **Worry subcluster (61)** | [32] | 0.13 (0.06) | 0.020 | 0.17 (0.05) | **$8.06 \times 10^{-4}$** | 0.04 (0.26) | 0.890 | 0.702 |
| **Number of sexual partners (64)** | [33] | 0.31 (0.04) | **$3.27 \times 10^{-12}$** | 0.36 (0.05) | **$9.00 \times 10^{-16}$** | 0.51 (0.20) | 0.011 | 0.309 |
| General risk tolerance (64) | [33] | 0.26 (0.06) | **$7.37 \times 10^{-6}$** | 0.28 (0.07) | **$5.93 \times 10^{-5}$** | 0.88 (0.25) | **$3.69 \times 10^{-4}$** | $9.62 \times 10^{-3}$ |
| **Insomnia (159)** | [34] | 0.05 (0.01) | **$1.90 \times 10^{-5}$** | 0.03 (0.01) | $5.31 \times 10^{-3}$ | 0.00 (0.05) | 0.993 | 0.288 |
| **Cognitive performance (134)** | [35] | -0.08 (0.02) | **$1.03 \times 10^{-3}$** | -0.05 (0.03) | 0.044 | -0.21 (0.12) | 0.086 | 0.282 |
| **Educational attainment (570)** | [35] | -0.22 (0.02) | **$1.32 \times 10^{-25}$** | -0.21 (0.02) | **$1.45 \times 10^{-17}$** | -0.24 (0.08) | $2.21 \times 10^{-3}$ | 0.781 |

P-values labeled in bold are significant after multiple testing correction. Traits labeled in bold are those having a causal effect on AUD risk by at least one method without evidencing horizontal pleiotropy (MR-Egger intercept $p > 0.05$ ). IVW: inverse-variance weighted (IVW) linear regression. DrnkWk: drinks per week. MDD: major depressive disorder. Depressed affect subcluster: depressed affect subcluster of neuroticism. Worry subcluster: worry subcluster of neuroticism.

### Joint Analysis of PAU and DrnkWk Using MTAG

We conducted a joint analysis of PAU and DrnkWk using MTAG, which can increase the power for each trait without introducing bias from sample overlap [10]. MTAG analysis increased the GWAS-equivalent sample size (*n*_Eq_) for PAU to 514,790, i.e., a 71.1% increase from the original effective sample size (*n*_E_ = 300,789, *n* = 435,563). In this analysis, we observed an increase in the number of independent variants for PAU to 119, 76 of which were conditionally independent (Supplementary Figure 9, Supplementary Table 14). For DrnkWk, the MTAG analysis increased the *n*_Eq_ to 612,968 from 537,352, which yielded 141 independent variants, 86 of which were conditionally independent (Supplementary Figure 10, Supplementary Table 15). MTAG analysis increased the observed *h*^2^ of PAU to 0.113 (se = 0.005) from 0.068 (se = 0.004) and of DrnkWk to 0.063 (se = 0.003) from the reported value of 0.042 (se = 0.002, Figure 2) [9].

The MTAG analysis also increased the power for the functional enrichment analysis. MAGMA gene set analysis for PAU after MTAG analysis detected 10 enriched Gene Ontology terms, including ‘regulation of nervous system development’ (p_Bonferroni_ = 8.80 × 10^−4^), ‘neurogenesis’ (p_Bonferroni_ = 0.010), and ‘synapse’ (p_Bonferroni_ = 0.046) (Supplementary Table 16).

## Discussion

We report here a genome-wide meta-analysis of PAU in 435,563 individuals of European ancestry from the MVP, PGC, and UKB datasets. MVP is a mega-biobank that has enrolled >750,000 subjects (for whom genotype data on 313,977 subjects was used in this study), with rich phenotype data assessed by questionnaires and from the EHR. Currently, MVP is the largest single cohort available with diagnostic information on AUD [3, 6]. PGC is a collaborative consortium that has led the effort to collect smaller cohorts with DSM-IV AD [2]. UKB is a population-level cohort with the largest available sample with AUDIT-P data [4].

Our discovery meta-analysis of PAU yielded 29 independent variants, of which 19 were novel, with 0.059 to 0.113 of the phenotypic variance explained in different cohorts or meta-analyses. The *h*^2^ in the Phase1-Phase2 MVP meta-analysis was 0.095 (se = 0.006), which was higher than MVP phase1: 0.056 (se = 0.004, in MVP phase1 where only the actual (as opposed to effective) sample size was used) [3]. The *h*^2^ of AD in PGC was 0.098 (se = 0.018), comparable to the reported liability-scale *h*^2^ (0.090, se = 0.019) [2]. Functional and heritability analyses consistently showed enrichments in brain regions and gene expression regulatory regions, providing biological insights into the etiology of PAU. Variation associated with gene expression in the brain is central to PAU risk, a conclusion that is also consistent with our previous GWASs in MVP of both alcohol consumption and AUD diagnosis [3]. The enrichments in regulatory regions point to specific brain tissues relevant to the causative genes; the specific interactions between 16 genes and 325 drugs may provide targets for the development of medications to manage PAU. Potential targets identified include the D_2_ dopamine receptor (encoded by *DRD2*) and phosphodiesterase 4B (encoded by *PDE4B*). The presence of risk variation at these loci also suggests the possibility that they may be “personalized medicine” targets as well.

We also found that PAU was significantly genetically correlated with 138 other traits. The top correlations were with substance use and substance-related disorders, MDD, schizophrenia, and several other neuropsychiatric traits. In a conceptually similar analysis, we performed a PheWAS of PAU PRS in BioVU, which confirmed the genetic correlations between PAU and multiple substance use disorders, mood disorders, and other psychiatric traits in an independent sample. We also used MR to infer causal effects of the above traits on liability to AUD (we tested AUD excluding UKB samples to avoid sample overlap) using selected genetic instruments. We found evidence of causal relationships from DrnkWk, ever smoked regularly, MDD, depressed affect subcluster, worry subcluster, number of sexual partners, insomnia, cognitive performance, and educational attainment to AUD risk, while cognitive performance and educational attainment showed protective effects on liability to AUD. For some of these observed effects, such as with schizophrenia, neuroticism, and general risk tolerance, we cannot exclude horizontal pleiotropy among our instrument variables. We could not test the reverse causality of AUD liability on other traits in the absence of large samples for those targeted traits, which are required to draw causal inferences. Thus we cannot rule out the possibility of bidirectional effects, which are plausible for several of these traits (e.g., MDD).

The study has other limitations. First, only European populations were included; therefore, the genetic architecture of PAU in other populations remains largely unknown. To date, the largest non-European sample to undergo GWAS for alcohol-related traits is African American (AA), which was reported in the MVP phase1 sample (17,267 cases; 39,381 controls, effective samples size 48,015), with the only associations detected being on chromosome 4 in the ADH gene locus (where several ADH genes map) [3]. Collection of substantial numbers of non-European subjects requires a concerted effort from our research field. Second, despite the high genetic correlation between AUD and AUDIT-P, they are not identical traits. We conducted a meta-analysis of the two traits to increase the power for the association study of PAU, consequently, associations specific to AUD or AUDIT-P could have been attenuated. Third, there was no opportunity for replication of the individual novel variants. Because the variants were detected in more than 430,000 subjects and have small effect sizes, a replication sample with adequate power would also have to be very large, and no such sample is currently available. To validate the findings, we conducted PRS analyses in three independent cohorts, which showed strong association with AUD. Although this indicates that our study had adequate power for variant detection, it does not address the validity of the individual variants discovered.

This is the largest GWAS study of PAU so far. Previous work has shown that the genetic architecture of AUD (and PAU) differs substantially from that of alcohol consumption [2–4]. There have been larger studies of alcohol quantity-frequency measures [9, 36]; alcohol consumption data are available in many EHRs, thus they were included in many studies of other primary traits, like cardiac disease. AUD diagnoses are collected much less commonly. The 3-item AUDIT-C is a widely-used measure of alcohol consumption often available in EHRs, but the full 10-item AUDIT, which allows the assessment of AUDIT-P, is not as widely available. Despite the high genetic correlation between, for example, PAU and DrnkWk (*r*_g_=0.77), very different patterns of genetic correlation and pleiotropy have been observed via LDSC and other methods for these different kinds of indices of alcohol use [2–5]. PAU captures pathological alcohol use: physiological dependence and/or significant medical consequences. Quantity/frequency measures may capture alcohol use that is in the normal, or anyway nonpathological, range. As such, we argue that although quantity/frequency measures are important for understanding the biology of habitual alcohol use, PAU is the more important, and more clearly pathological, trait. These circumstances underscore the importance of assembling a large GWAS sample of PAU to inform the biology of PAU, and our study moves towards this goal via the identification of numerous previously-unidentified risk loci: we increased known PAU loci from 10 to 29, nearly tripling our knowledge of specific risk regions. Similarly, we identified 66 gene-based associations, of which 46 were novel – again roughly tripling current knowledge. MTAG analysis increased locus discovery to 119, representing 76 independent loci, by levering information from DrnkWk [9]. By the same token, we provide a major increment in information about the biology of PAU, providing considerable fodder for future in-vitro and animal studies, which will be required to delineate the biology and function associated with each risk variant. We anticipate that this knowledge may lead to improvements in treatment and treatment personalization, a major ultimate goal of the work.

## Methods

### MVP datasets

The MVP is a mega-biobank supported by the U.S. Department of Veterans Affairs (VA), enrollment for which began in 2011 and is ongoing. Phenotypic data were collected using questionnaires and the VA electronic health records (EHR), and a blood sample was obtained from each participant for genetic studies. Two phases of genotypic data have been released and were included in this study. MVP phase1 contains 353,948 subjects, of whom 202,004 European Americans (EA) with AUD diagnoses were included in a previous GWAS and the summary statistics were used in this study [3]. MVP phase2 released data on another 108,416 subjects, of whom 65,387 EAs with AUD diagnosis information were included in this study. Following the same procedures as for MVP phase1, participants with at least one inpatient or two outpatient alcohol-related ICD-9/10 codes from 2000 to 2018 were assigned a diagnosis of AUD.

Ethics statement: The Central VA Institutional Review Board (IRB) and site-specific IRBs approved the MVP study. All relevant ethical regulations for work with human subjects were followed in the conduct of the study and informed consent was obtained from all participants.

Genotyping for both phases of MVP was performed using a customized Affymetrix Biobank Array. Imputation and quality control methods for MVP phase1 were described in detail in Kranzler et al. [3]. Similar methods were used for MVP phase2. Before imputation, phase2 subjects or SNPs with genotype call rate < 0.9 or high heterozygosity were removed, leaving 108,416 subjects and 668,324 SNPs. Imputation for MVP phase2 was done separately from phase1; both were performed with EAGLE2 [37] and Minimac3 [38] using 1000 Genomes Project phase 3 data [39] as the reference panel. Imputed genotypes with posterior probability ≥ 0.9 were transferred to best guess genotypes (the rest were treated as missing genotype calls). A total of 6,635,093 SNPs with INFO scores > 0.7, genotype call rates or best guess rates > 0.95, Hardy Weinberg equilibrium p value < 1 × 10^−6^, minor allele frequency (MAF) > 0.001 were remained for GWAS.

We removed subjects with mismatched genotypic and phenotypic sex and one subject randomly from each pair of related individuals (kinship coefficient threshold = 0.0884), leaving 107,438 phase2 subjects for subsequent analyses. We used the same processes as MVP phase1 to define EAs. First, we ran principal components analysis (PCA) on 74,827 common SNPs (MAF > 0.05) shared by MVP and the 1000 Genomes phase 3 reference panels using FastPCA [40]. Then we clustered each participant into the nearest reference population according to the Euclidean distances between the participant and the centers of the 5 reference populations using the first 10 PCs. A second PCA was performed for participants who were clustered to the reference European population (EUR), and outliers were removed if any of the first 10 PCs were > 3 standard deviations from the mean, leaving 67,268 EA subjects.

Individuals < 22 or > 90 years of age and those with a missing AUD diagnosis were removed from the analyses, leaving 65,387 phase2 EAs (11,337 cases; 54,050 controls). GWAS was then performed on the MVP phase2 dataset. We used logistic regression implemented in PLINK v1.90b4.4 [41] for the AUD GWAS correcting for age, sex, and the first 10 PCs.

### PGC summary statistics

We used the 46,568 European ancestry subjects (11,569 cases and 34,999 controls) from 27 cohorts that were analyzed by the Psychiatric Genomics Consortium (PGC). The phenotype was lifetime DSM-IV diagnosis of alcohol dependence (AD). The summary data were downloaded from the PGC website (https://www.med.unc.edu/pgc/) with full agreement to the PGC conditions. Allele frequencies were not reported in the summary data. We used allele frequencies from the 1000 Genome European sample as proxy measures in PGC for some downstream analyses.

### UK Biobank summary statistics

The UK Biobank (UKB) included 121,604 White-British unrelated subjects with available AUDIT-P scores. Past-year AUDIT-P was assessed by 7 questions: 1). Frequency of inability to cease drinking; 2). Frequency of failure to fulfil normal expectations due to drinking alcohol; 3). Frequency of needing morning drink of alcohol after heavy drinking session; 4). Frequency of feeling guilt or remorse after drinking alcohol; 5). Frequency of memory loss due to drinking alcohol; 6). Ever been injured or injured someone else through drinking alcohol; 7). Ever had known person concerned about, or recommend reduction of, alcohol consumption. The AUDIT-P was log_10_-transformed for GWAS (see ref [4] for details). We removed SNPs with INFO < 0.7 or call rate < 0.95.

### Meta-analyses

Meta-analyses were performed using METAL [42]. The meta-analysis within MVP (for the purpose of genetic correlation analysis with PGC AD) was conducted using an inverse variance weighted method because the two subsets were from the same cohort. The meta-analyses for AUD (MVP+PGC) and PAU (MVP+PGC+UKB) were performed using the sample size weighted method. Given the unbalanced ratios of cases to controls in MVP samples, we calculated effective sample sizes for meta-analysis following the approach used by the PGC:

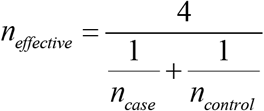

The calculated effective sample sizes in MVP and reported effective sample sizes in PGC were used in meta-analyses and all downstream analyses. AUDIT-P in UKB is a continuous trait, so we used actual sample sizes for that trait. For the AUD meta-analysis, variants present in only one sample (except MVP phase1 which is much larger than the others) or with heterogeneity test p-value < 5 × 10^−8^ were removed, leaving 7,003,540 variants. For the PAU meta-analysis, variants present in only one sample (except MVP phase1 or UKB) or with heterogeneity test p-value < 5 × 10^−8^ and variants with effective sample size < 45,118 (15% of the total effective sample size) were removed, leaving 14,069,427 variants.

### AUD polygenic risk score in UKB

We calculated AUD polygenic risk scores (PRS) for each of the 82,930 unrelated subjects in UKB who had AUDIT-P information [7]. A PRS was calculated as the sum of the number of effective alleles with p-values less than a given threshold, weighted by the effect sizes from AUD meta-analysis (MVP+PGC). We analyzed 10 p-value thresholds: 5 × 10^−8^, 1 × 10^−7^, 1 × 10^−6^, 1 × 10^−5^, 1 × 10^−4^, 0.001, 0.05, 0.3, 0.5, and 1, and clumped the AUD summary data by LD with *r*^2^ < 0.3 in a 500 kb window. Then we tested the association between AUD PRS and AUDIT-P, corrected for age, sex, and 10 PCs. The analysis was performed using PRSice-2 [43].

### Independent variants and conditional analyses

We identified the independent variant (p < 5 × 10^−8^) in each locus (1 Mb genomic window) based on the smallest p value and *r*^2^ < 0.1 with other independent variants. Variants with p < 1 × 10^−8^ and *r*^2^ > 0.1 with respect to the independent variants were assigned to the independent variant’s clump. Any two independent variants less than 1 Mb apart whose clumped regions overlapped were merged into one locus. Given the known long-range LD for the ADH gene cluster on chromosome 4, we defined chr4q23–q24 (~97.2 Mb – 102.6 Mb) as one locus. When multiple independent variants were present in a locus, we ran conditional analyses using GCTA-COJO [44] to define conditionally independent variants. For each variant other than the most significant one (index), we tested the marginal associations conditioning on the index variant using Europeans (n = 503) from the 1000 Genomes as the LD reference sample. Variants with significant marginal associations (p < 5 × 10^−8^) were defined as conditionally independent variants (i.e., independent when conditioned on other variants in the region) and subject to another round of conditional analyses for each significant association.

For the conditionally independent variants for AUD or PAU, we also conducted a multi-trait analysis conditioning on GSCAN drinks per week [9] using GCTA-mtCOJO [45] to identify variants associated with AUD or PAU, but not drinks per week, i.e., not alcohol consumption alone. Europeans from the 1000 Genomes were used as the LD reference. For variants missing in GSCAN, we used proxy variants (p < 5 × 10^−8^) in high LD with the locus for analyses. Whereas conditional analyses require the beta (effect size) and standard error, we calculated these using Z-scores (*z*), allele frequency (*p*) and sample size (*n*) from the meta-analyses [46]:

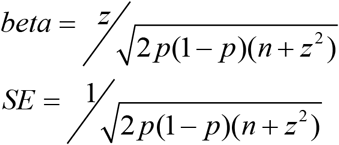

### Gene-based association analysis

Gene-based association analysis for PAU was performed using MAGMA implemented in FUMA [17, 18], which uses a multiple regression approach to detect multi-marker effects that account for SNP p-values and LD between markers. We used default settings to analyze 18,952 autosomal genes, with p < 2.64 × 10^−6^ (0.05/18,952) considered GWS.

### Drug-gene interaction

For the genes identified as significant by MAGMA, we examined drug-gene interaction through Drug Gene Interaction Database (DGIdb) v3.0.2 [11] (http://www.dgidb.org/), a database integrated drug–gene interaction information resource based on 30 sources.

### SNP-based *h*^2^ and partitioning heritability enrichment

LDSC [12] was used to estimate the SNP-based *h*^2^ for common SNPs mapped to HapMap3 [47], using Europeans from the 1000 Genomes Project [39] as the LD reference panel. We excluded the major histocompatibility complex (MHC) region (chr6: 26–34Mb).

We conducted portioning *h*^2^ enrichment analyses for PAU using LDSC in different models [13, 14]. First, a baseline model consisting of 52 functional categories was analyzed, which included genomic features (coding, intron, UTR etc), regulatory annotations (promoter, enhancer etc), epigenomic annotations (H3K27ac, H3K4me1, H3K3me3 etc) and others (see ref [13] for details, Supplementary Figure 5). We then analyzed cell type group *h*^2^ enrichments with 10 cell types: central nervous system (CNS), adrenal and pancreas, immune and hematopoietic, skeletal muscle, gastrointestinal, liver, cardiovascular, connective tissue and bone, kidney, and other (see ref [13] for details, Supplementary Figure 4). Third, we used LDSC to test for enriched heritability in regions surrounding genes with the highest tissue-specific expression using 53 human tissue or cell type RNA-seq data from the Genotype-Tissue Expression Project (GTEx) [16], or enriched heritability in epigenetic markers from 396 human epigenetic annotations (six features in a subset of 88 primary cell types or tissues) from the Roadmap Epigenomics Consortium [15] (see ref [14] for details, Supplementary Figure 6, Supplementary Table 6). For each model, the number of tested annotations was used to calculate a Bonferroni corrected p-value < 0.05 as a significance threshold.

### Gene-set and functional enrichment

We performed gene-set analysis for PAU for curated gene sets and Gene Ontology (GO) terms using MAGMA [17, 18]. We then used MAGMA for gene-property analyses to test the relationships between tissue-specific gene expression profiles and PAU-gene associations. We analyzed gene expression data from 53 GTEx (v7) tissues. We also performed gene-set analysis on the 152 prioritized genes using MAGMA. Gene sets with adjusted p-value < 0.05 were considered as significant.

### Genetic correlation

We estimated the genetic correlation (*r*_g_) between traits using LDSC [48]. For PAU, we estimated the *r*_g_ with 218 published traits in LD Hub [49], 487 unpublished traits from the UK Biobank (integrated in LD Hub), and recently published psychiatric and behavioral traits [9, 30, 32–35, 50–54], bringing the total number of tested traits to 715 (Supplementary Table 8). For traits reported in multiple studies or in UKB, we selected the published version of the phenotype or used the largest sample size. Bonferroni correction was applied and correlation was considered significant at a p-value threshold of 6.99 × 10^−5^.

### S-PrediXcan and S-MultiXcan

To perform transcriptome-wide association analysis, we used S-PrediXcan [23] (a version of PrediXcan that uses GWAS summary statistics [55]) to integrate transcriptomic data from GTEx [16] and the Depression Genes and Networks study (DGN) [24] to analyze the summary data from the PAU meta-analysis. Forty-eight tissues with sample size > 70 from GTEx release v7 were analyzed, totaling 10,294 samples. DGN contains RNA sequencing data from whole blood of 992 genotyped individuals. The transcriptome prediction model database and the covariance matrices of the SNPs within each gene model were downloaded from the PredictDB repository (http://predictdb.org/,2018-01-08 release). Only individuals of European ancestry in GTEx were analyzed. S-PrediXcan was performed for each of the 49 tissues (48 from GTEx and 1 from DGN), for a total of 254,345 gene-tissue pairs. Significant association was determined by Bonferroni correction (p < 1.97 × 10^−7^).

Considering the limited eQTL sample size for any single tissue and the substantial sharing of eQTLs across tissues, we applied S-MultiXcan [26], which integrates evidence across multiple tissues using multivariate regression to improve association detection. Forty-eight tissues from GTEx were analyzed jointly. The threshold for condition number of eigenvalues was set to 30 when truncating singular value decomposition (SVD) components. In total, 25,626 genes were tested in S-MultiXcan, leading to a significant p-value threshold of 1.95 × 10^−6^ (0.05/25,626).

### PAU PRS for phenome-wide associations

Polygenic scores were generated using PRS-CS [56] on all genotyped individuals of European descent (n = 67,588) in Vanderbilt University Medical Center’s EHR-linked biobank, BioVU. PRS-CS uses a Bayesian framework to model linkage disequilibrium from an external reference set and a continuous shrinkage prior on SNP effect sizes. We used 1000 Genomes Project Phase 3 European sample [39] as the LD reference. Additionally, we used the PRS-CS-auto option, which allows the software to learn the continuous shrinkage prior from the data. Polygenic scores were constructed from PRS-CS-auto adjusted summary statistics containing 811,292 SNPs. All individuals used for polygenic scoring were genotyped on the Illumina Multi-Ethnic Global Array (MEGA). Genotypes were filtered for SNP (95%) and individual (98%) call rates, sex discrepancies, and excessive heterozygosity. For related individuals, one of each pair was randomly removed (pi_hat > 0.2). SNPs showing significant associations with genotyping batch were removed. Genetic ancestry was determined by principal component analysis performed using EIGENSTRAT [57]. Imputation was completed using the Michigan Imputation Server [38] and the Haplotype Reference Consortium [58] as the reference panel. Genotypes were then converted to hard calls, and filtered for SNP imputation quality (R^2^ > 0.3), individual missingness (>2%), SNP missingness (>2%), minor allele frequency (<1%) and Hardy-Weinberg Equilibrium (p > 1 × 10^−10^). The resulting dataset contained 9,330,483 SNPs on 67,588 individuals of European ancestry.

We conducted a phenome-wide association study (PheWAS) [59] of the PAU PRS, by fitting a logistic regression model to 1,372 case/control phenotypes to estimate the odds of each diagnosis given the PAU polygenic score, controlling for sex, median age across the medical record, top 10 principal components of ancestry, and genotyping batch. We required the presence of at least two International Classification of Disease (ICD) codes that mapped to a PheWAS disease category (Phecode Map 1.2) to assign “case” status. A phenotype was required to have at least 100 cases to be included in the analysis. PheWAS analyses were run using the PheWAS R package [60]. Bonferroni correction was applied to test for significance (p < 0.05/1,372).

### PAU PRS in independent samples

We calculated PAU PRS in three independent samples, where we tested the association between PAU PRS and AD, corrected for age, sex, and 10 PCs. Ten p-value thresholds were applied in all samples.

#### iPSYCH Group

DNA samples for cases and controls were obtained from newborn bloodspots linked to population registry data [61]. Cases were identified with the ICD-10 code F10.2 (AD; *n* = 944); controls were from the iPSYCH group (*n* = 11,408; *n*_effective_ = 3,487)). The iPSYCH sample was genotyped on the Psych Array (Illumina, San Diego, CA, US). GWAS QC, imputation against the 1,000 Genomes Project panel [39] and association analysis using the Ricopili pipeline [62] were performed.

#### UCL Psych Array

Cases were identified with ICD-10 code F10.2 (*n* = 1,698) and comprised 492 individuals with a diagnosis of alcoholic hepatitis who had participated in the STOPAH (Steroids or Pentoxifylline for Alcoholic Hepatitis) trial (ISRCTN88782125; EudraCT Number: 2009-013897-42) and 1,206 subjects recruited from the AD arm of the DNA Polymorphisms in Mental Health (DPIM) study; controls were UK subjects who had either been screened for an absence of mental illness and harmful substance use (*n* = 776), or were random blood donors (n-452; total *n* = 1,228; *n*_effective_ = 2,851). The sample was genotyped on the Psych Array (Illumina, San Diego, CA, US). GWAS QC was performed using standard methods and imputation was done using the haplotype reference consortium (HRC) panel [63] on the Sanger Imputation server (https://imputation.sanger.ac.uk/). Association testing was performed using Plink1.9 [41].

#### UCL Core Exome Array

Cases had an ICD-10 diagnosis of F10.2 (*n* = 637), including 324 individuals with a diagnosis of alcoholic hepatitis who had participated in the STOPAH trial and 313 subjects recruited from the AD arm of the DPIM study; controls were unrelated UK subjects from the UK Household Longitudinal Study (UKHLS; *n* = 9,189; *n*_effective_ = 2,383). The sample was genotyped on the Illumina Human Core Exome Array (Illumina, San Diego, CA, US). GWAS QC was performed using standard methods and imputation was done using the HRC panel [63] on the Sanger Imputation server (https://imputation.sanger.ac.uk/). Association testing was performed with Plink1.9 [41].

### Mendelian Randomization

We used Mendelian Randomization (MR) to investigate the causal relationships with PAU liability of the many traits that were significantly genetically correlated (p < 6.99 × 10^−5^). However, all or most of the published traits in recent large GWAS include UKB data. To avoid biases caused by overlapping samples in MR analysis, we only tested the relationship between published traits and AUD (MVP+PGC). For robust causal effect inference, we limited the traits studied to those with more than 30 available instruments (association p < 5 × 10^−8^). Only the causal effects of liability to other exposures on AUD risk were tested given that there are only 24 independent variants for AUD. In total, 13 exposures were analyzed (Table 2).

Three methods, weighted median [28], inverse-variance weighted (IVW, random-effects model) [27], and MR-Egger [29], implemented in the R package “MendelianRandomization v0.3.0” [64] were used for MR inference. Evidence of pleiotropic effects was examined by the MR-Egger intercept test, where a non-zero intercept indicates directional pleiotropy [29]. Instrumental variants that are associated with PAU (p < 5 × 10^−8^) were removed. For instrumental variants missing in the PAU summary data, we used the results of the best-proxy variant with the highest LD (*r*^2^ > 0.8) with the missing variant. If the MAF of the missing variant was < 0.01, or none of the variants within 200 kb had LD *r*^2^ > 0.8, we removed the instrumental variant from the analysis.

### MTAG between PAU and drinks per week

Multiple trait analysis between PAU and drinks per week (DrnkWk) from GSCAN was performed on summary statistics with multi-trait analysis of GWAS (MTAG) v1.0.7 [10]. The summary data of DrnkWk were generated from 537,352 subjects, excluding the 23andMe samples that were not available to us for inclusion. We analyzed variants with a minimum effective sample size of 80,603 (15%) in DrnkWk and a minimum effective sample size of 45,118 (15%) in PAU, which left 10,613,246 overlapping variants.

## Acknowledgements

This research used data from the Million Veteran Program, Office of Research and Development, Veterans Health Administration, and was supported by award #1I01BX003341 and CSP575B. This publication does not represent the views of the Department of Veterans Affairs or the United States Government. Supported also by NIH (NIAAA) P50 AA12870, a NARSAD Young Investigator Grant from the Brain & Behavior Research Foundation (HZ), and NIH grants 5T32GM080178 (JMS) and K02DA32573 (AA); and the NIHR Imperial Biomedical Research Centre (SRA and MRT). This research also used summary data from the Psychiatric Genomics Consortium (PGC) Substance Use Disorders (SUD) working group. The PGC-SUD is supported by funds from NIDA and NIMH to MH109532 and, previously, had analyst support from NIAAA to U01AA008401 (COGA). PGC-SUD gratefully acknowledges its contributing studies and the participants in those studies, without whom this effort would not be possible. This research also used summary data from UK Biobank, a population-based sample of participants whose contributions we gratefully acknowledge. We thank the iPSYCH-Broad Consortium for access to data on the iPSYCH cohort. The iPSYCH project is funded by the Lundbeck Foundation (R102-A9118 and R155-2014-1724) and the universities and university hospitals of Aarhus and Copenhagen. Genotyping of iPSYCH samples was supported by grants from the Lundbeck Foundation and the Stanley Foundation, The Danish National Biobank resource was supported by the Novo Nordisk Foundation. Data handling and analysis on the GenomeDK HPC facility was supported by NIMH (1U01MH109514-01 to ADB). High-performance computer capacity for handling and statistical analysis of iPSYCH data on the GenomeDK HPC facility was provided by the Centre for Integrative Sequencing, iSEQ, Aarhus University, Denmark (grant to ADB).

## References

1. GBD1 2016 Alcohol Collaborators., Alcohol use and burden for 195 countries and territories, 1990-2016: a systematic analysis for the Global Burden of Disease Study 2016. Lancet, 2018. 392(10152): p. 1015–1035.

2. Walters, R.K., et al., Transancestral GWAS of alcohol dependence reveals common genetic underpinnings with psychiatric disorders. Nat Neurosci, 2018. 21(12): p. 1656–1669.

3. Kranzler, H.R., et al., Genome-wide association study of alcohol consumption and use disorder in 274,424 individuals from multiple populations. Nat Commun, 2019. 10(1): p. 1499.

4. Sanchez-Roige, S., et al., Genome-Wide Association Study Meta-Analysis of the Alcohol Use Disorders Identification Test (AUDIT) in Two Population-Based Cohorts. Am J Psychiatry, 2019. 176(2): p. 107–118.

5. Gelernter, J., et al., Genome-wide Association Study of Maximum Habitual Alcohol Intake in >140,000 U.S. European and African American Veterans Yields Novel Risk Loci. Biol Psychiatry, 2019.

6. Gaziano, J.M., et al., Million Veteran Program: A mega-biobank to study genetic influences on health and disease. J Clin Epidemiol, 2016. 70: p. 214–23.

7. Bycroft, C., et al., The UK Biobank resource with deep phenotyping and genomic data. Nature, 2018. 562(7726): p. 203–209.

8. Gelernter, J., et al., Genome-wide association study of alcohol dependence:significant findings in African- and European-Americans including novel risk loci. Mol Psychiatry, 2014. 19(1): p. 41–9.

9. Liu, M., et al., Association studies of up to 1.2 million individuals yield new insights into the genetic etiology of tobacco and alcohol use. Nat Genet, 2019. 51(2): p. 237–244.

10. Turley, P., et al., Multi-trait analysis of genome-wide association summary statistics using MTAG. Nat Genet, 2018. 50(2): p. 229–237.

11. Cotto, K.C., et al., DGIdb 3.0: a redesign and expansion of the drug-gene interaction database. Nucleic Acids Res, 2018. 46(D1): p. D1068–D1073.

12. Bulik-Sullivan, B.K., et al., LD Score regression distinguishes confounding from polygenicity in genome-wide association studies. Nat Genet, 2015. 47(3): p. 291–5.

13. Finucane, H.K., et al., Partitioning heritability by functional annotation using genome-wide association summary statistics. Nat Genet, 2015. 47(11): p. 1228–35.

14. Finucane, H.K., et al., Heritability enrichment of specifically expressed genes identifies disease-relevant tissues and cell types. Nat Genet, 2018. 50(4): p. 621–629.

15. Roadmap Epigenomics Consortium, et al., Integrative analysis of 111 reference human epigenomes. Nature, 2015. 518(7539): p. 317–30.

16. GTEx Consortium, Genetic effects on gene expression across human tissues. Nature, 2017. 550(7675): p. 204–213.

17. Watanabe, K., et al., Functional mapping and annotation of genetic associations with FUMA. Nat Commun, 2017. 8(1): p. 1826.

18. de Leeuw, C.A., et al., MAGMA: generalized gene-set analysis of GWAS data. PLoS Comput Biol, 2015. 11(4): p. e1004219.

19. Marees, A.T., et al., Potential influence of socioeconomic status on genetic correlations between alcohol consumption measures and mental health. Psychol Med, 2019: p. 1–15.

20. Grant, B.F., et al., Epidemiology of DSM-5 Alcohol Use Disorder: Results From the National Epidemiologic Survey on Alcohol and Related Conditions III. JAMA Psychiatry, 2015. 72(8): p. 757–66.

21. Andersen, A.M., et al., Polygenic Scores for Major Depressive Disorder and Risk of Alcohol Dependence. JAMA Psychiatry, 2017. 74(11): p. 1153–1160.

22. Zhou, H., et al., Genetic Risk Variants Associated With Comorbid Alcohol Dependence and Major Depression. JAMA Psychiatry, 2017. 74(12): p. 1234–1241.

23. Barbeira, A.N., et al., Exploring the phenotypic consequences of tissue specific gene expression variation inferred from GWAS summary statistics. Nat Commun, 2018. 9(1): p. 1825.

24. Battle, A., et al., Characterizing the genetic basis of transcriptome diversity through RNA-sequencing of 922 individuals. Genome Res, 2014. 24(1): p. 14–24.

25. Edenberg, H.J. and J.N. McClintick, Alcohol Dehydrogenases, Aldehyde Dehydrogenases, and Alcohol Use Disorders: A Critical Review. Alcohol Clin Exp Res, 2018. 42(12): p. 2281–2297.

26. Barbeira, A.N., et al., Integrating predicted transcriptome from multiple tissues improves association detection. PLoS Genet, 2019. 15(1): p. e1007889.

27. Bowden, J., et al., A framework for the investigation of pleiotropy in two-sample summary data Mendelian randomization. Stat Med, 2017. 36(11): p. 1783–1802.

28. Bowden, J., et al., Consistent Estimation in Mendelian Randomization with Some Invalid Instruments Using a Weighted Median Estimator. Genet Epidemiol, 2016. 40(4): p. 304–14.

29. Bowden, J., G. Davey Smith, and S. Burgess, Mendelian randomization with invalid instruments: effect estimation and bias detection through Egger regression. Int J Epidemiol, 2015. 44(2): p. 512–25.

30. Howard, D.M., et al., Genome-wide meta-analysis of depression identifies 102 independent variants and highlights the importance of the prefrontal brain regions. Nat Neurosci, 2019. 22(3): p. 343–352.

31. Schizophrenia Working Group of the Psychiatric Genomics, C., Biological insights from 108 schizophrenia-associated genetic loci. Nature, 2014. 511(7510): p. 421–7.

32. Nagel, M., et al., Meta-analysis of genome-wide association studies for neuroticism in 449,484 individuals identifies novel genetic loci and pathways. Nat Genet, 2018. 50(7): p. 920–927.

33. Karlsson Linner, R., et al., Genome-wide association analyses of risk tolerance and risky behaviors in over 1 million individuals identify hundreds of loci and shared genetic influences. Nat Genet, 2019. 51(2): p. 245–257.

34. Jansen, P.R., et al., Genome-wide analysis of insomnia in 1,331,010 individuals identifies new risk loci and functional pathways. Nat Genet, 2019. 51(3): p. 394–403.

35. Lee, J.J., et al., Gene discovery and polygenic prediction from a genome-wide association study of educational attainment in 1.1 million individuals. Nat Genet, 2018. 50(8): p. 1112–1121.

36. Evangelou, E., et al., New alcohol-related genes suggest shared genetic mechanisms with neuropsychiatric disorders. Nat Hum Behav, 2019.

37. Loh, P.R., et al., Reference-based phasing using the Haplotype Reference Consortium panel. Nat Genet, 2016. 48(11): p. 1443–1448.

38. Das, S., et al., Next-generation genotype imputation service and methods. Nat Genet, 2016. 48(10): p. 1284–1287.

39. 1000 Genomes Project Consortium, A global reference for human genetic variation. Nature, 2015. 526(7571): p. 68–74.

40. Galinsky, K.J., et al., Fast Principal-Component Analysis Reveals Convergent Evolution of ADH1B in Europe and East Asia. Am J Hum Genet, 2016. 98(3): p. 456–472.

41. Chang, C.C., et al., Second-generation PLINK: rising to the challenge of larger and richer datasets. Gigascience, 2015. 4: p. 7.

42. Willer, C.J., Y. Li, and G.R. Abecasis, METAL: fast and efficient meta-analysis of genomewide association scans. Bioinformatics, 2010. 26(17): p. 2190–1.

43. Euesden, J., C.M. Lewis, and P.F. O’Reilly, PRSice: Polygenic Risk Score software. Bioinformatics, 2015. 31(9): p. 1466–8.

44. Yang, J., et al., Conditional and joint multiple-SNP analysis of GWAS summary statistics identifies additional variants influencing complex traits. Nat Genet, 2012. 44(4): p. 369–75, S1–3.

45. Zhu, Z., et al., Causal associations between risk factors and common diseases inferred from GWAS summary data. Nat Commun, 2018. 9(1): p. 224.

46. Zhu, Z., et al., Integration of summary data from GWAS and eQTL studies predicts complex trait gene targets. Nat Genet, 2016. 48(5): p. 481–7.

47. International HapMap Consortium, et al., Integrating common and rare genetic variation in diverse human populations. Nature, 2010. 467(7311): p. 52–8.

48. Bulik-Sullivan, B., et al., An atlas of genetic correlations across human diseases and traits. Nat Genet, 2015. 47(11): p. 1236–41.

49. Zheng, J., et al., LD Hub: a centralized database and web interface to perform LD score regression that maximizes the potential of summary level GWAS data for SNP heritability and genetic correlation analysis. Bioinformatics, 2017. 33(2): p. 272–279.

50. Pasman, J.A., et al., GWAS of lifetime cannabis use reveals new risk loci, genetic overlap with psychiatric traits, and a causal influence of schizophrenia. Nat Neurosci, 2018. 21(9): p. 1161–1170.

51. Savage, J.E., et al., Genome-wide association meta-analysis in 269,867 individuals identifies new genetic and functional links to intelligence. Nat Genet, 2018. 50(7): p. 912–919.

52. Demontis, D., et al., Discovery of the first genome-wide significant risk loci for attention deficit/hyperactivity disorder. Nat Genet, 2019. 51(1): p. 63–75.

53. Jansen, I.E., et al., Genome-wide meta-analysis identifies new loci and functional pathways influencing Alzheimer’s disease risk. Nat Genet, 2019. 51(3): p. 404–413.

54. Stahl, E.A., et al., Genome-wide association study identifies 30 loci associated with bipolar disorder. Nat Genet, 2019. 51(5): p. 793–803.

55. Gamazon, E.R., et al., A gene-based association method for mapping traits using reference transcriptome data. Nat Genet, 2015. 47(9): p. 1091–8.

56. Ge, T., et al., Polygenic prediction via Bayesian regression and continuous shrinkage priors. Nat Commun, 2019. 10(1): p. 1776.

57. Price, A.L., et al., Principal components analysis corrects for stratification in genome-wide association studies. Nat Genet, 2006. 38(8): p. 904–9.

58. McCarthy, S., et al., A reference panel of 64,976 haplotypes for genotype imputation. Nat Genet, 2016. 48(10): p. 1279–83.

59. Denny, J.C., et al., Systematic comparison of phenome-wide association study of electronic medical record data and genome-wide association study data. Nat Biotechnol, 2013. 31(12): p. 1102–10.

60. Carroll, R.J., L. Bastarache, and J.C. Denny, R PheWAS: data analysis and plotting tools for phenome-wide association studies in the R environment. Bioinformatics, 2014. 30(16): p. 2375–6.

61. Pedersen, C.B., et al., The iPSYCH2012 case-cohort sample: new directions for unravelling genetic and environmental architectures of severe mental disorders. Mol Psychiatry, 2018. 23(1): p. 6–14.

62. Lam, M., et al., RICOPILI: Rapid Imputation for COnsortias PIpeLIne. BioRxiv, 2019. https://doi.org/10.1101/587196.

63. McCarthy, S., et al., A reference panel of 64,976 haplotypes for genotype imputation. Nat Genet, 2016. 48(10): p. 1279–83.

64. Yavorska, O.O. and S. Burgess, MendelianRandomization: an R package for performing Mendelian randomization analyses using summarized data. Int J Epidemiol, 2017. 46(6): p. 1734–1739.

