## Supplementary Figures for "Meta-analysis of problematic alcohol use in 435,563 individuals identifies 29 risk variants and yields insights into biology, pleiotropy and causality"

**Supplementary Figure 1. Manhattan and QQ plots for AUD meta-analysis (MVP+PGC).**

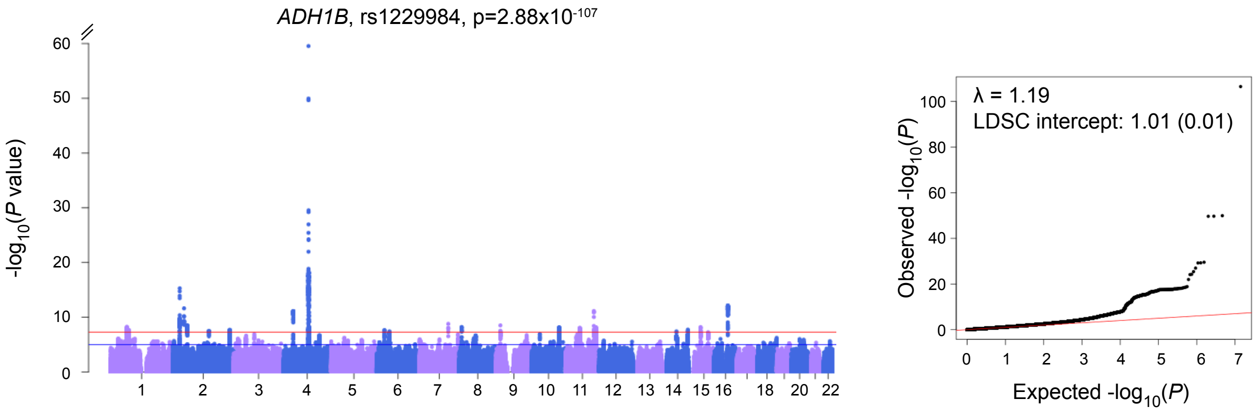

**Supplementary Figure 2. Quantiles for polygenic risk score of AUD meta-analysis associated with AUDIT-P in UKB.**

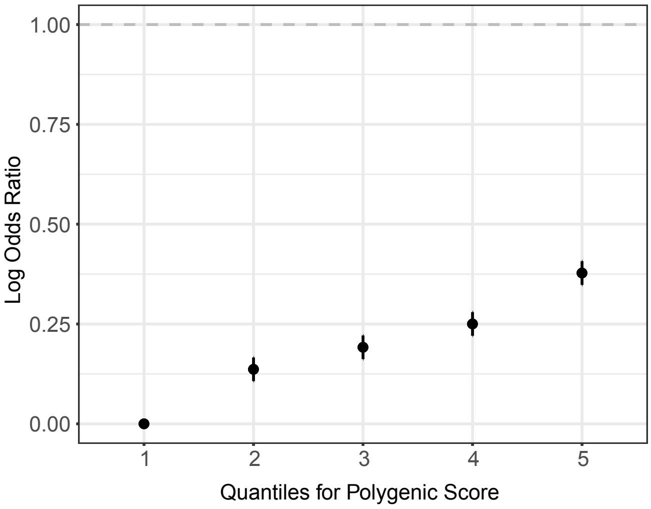

**Supplementary Figure 3. Manhattan and QQ plots for PAU.**

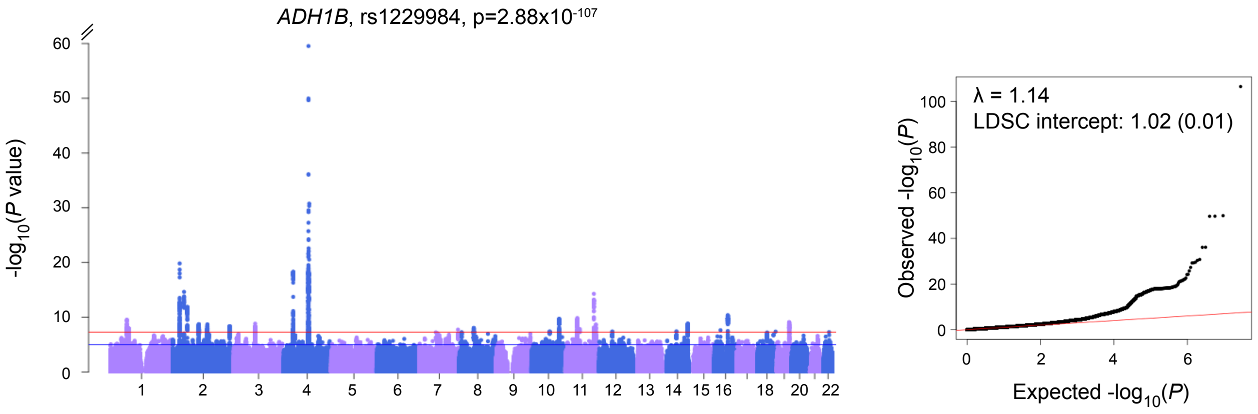

**Supplementary Figure 4. Cell type group partitioning heritability enrichment for PAU using LDSC.** The dashed line is the cutoff for Bonferroni-corrected significance. CNS: central nervous system.

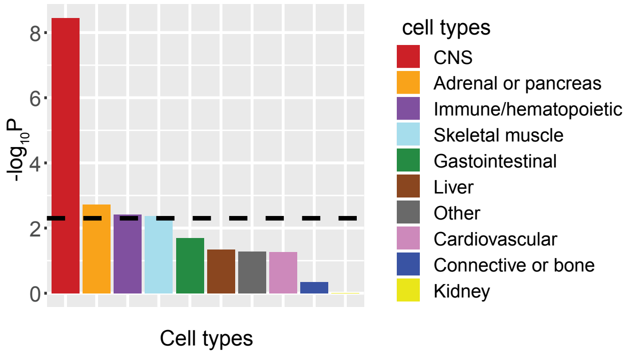

**Supplementary Figure 5. Functional category partitioning heritability enrichment for PAU using LDSC baseline model.** The categories labled by asterisks are significant after Bonferroni correction. For each of the 24 main annotations, there is an annotation extended 500 bp (ext) from the original one. There are 4 annotations with 100bp windows around the peaks (peaks) from the original ones.

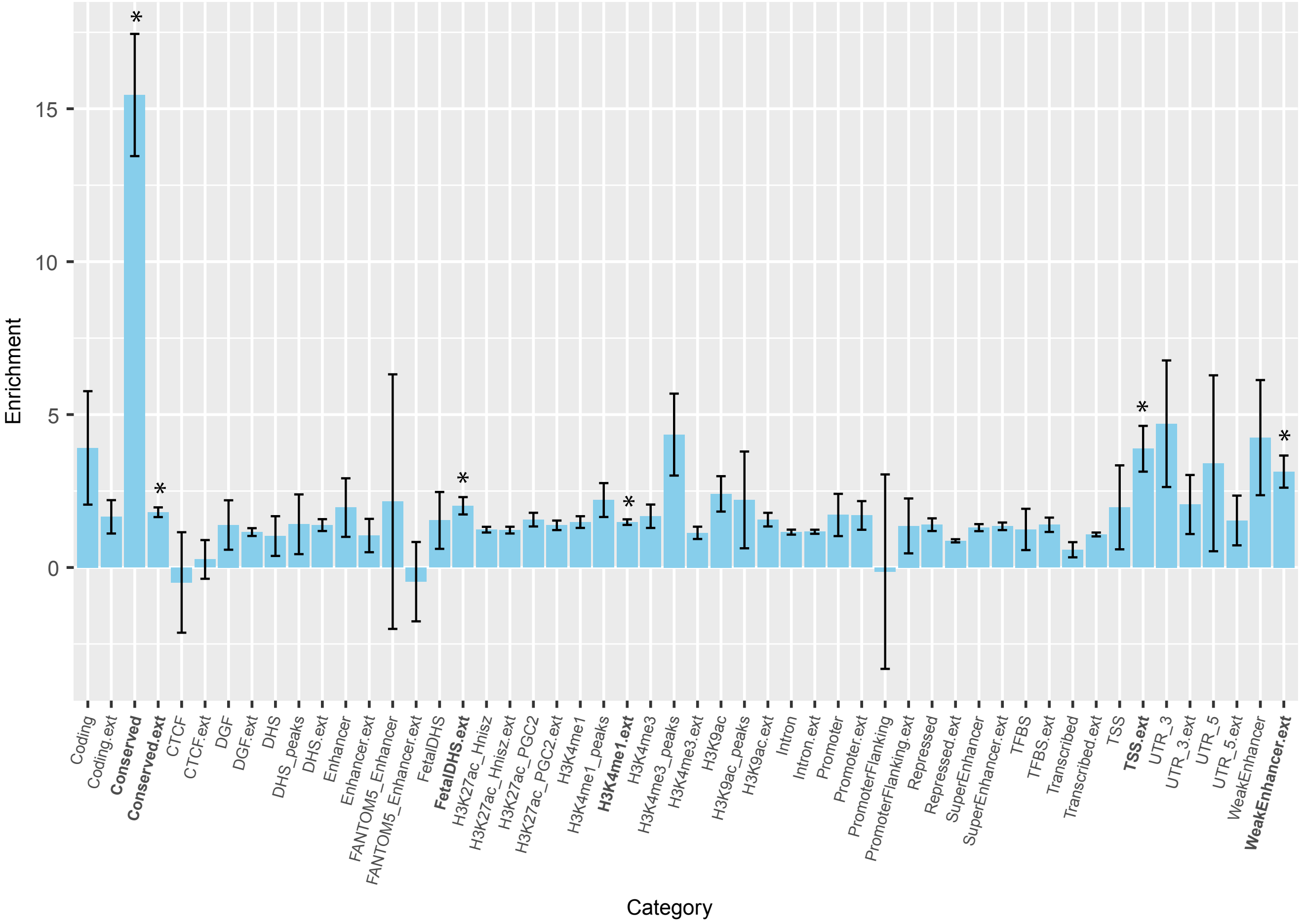

**Supplementary Figure 6. GTEx tissues or cell types partitioning heritability enrichment for PAU using LDSC.** The dashed line is the cutoff for Bonferroni-corrected significance.

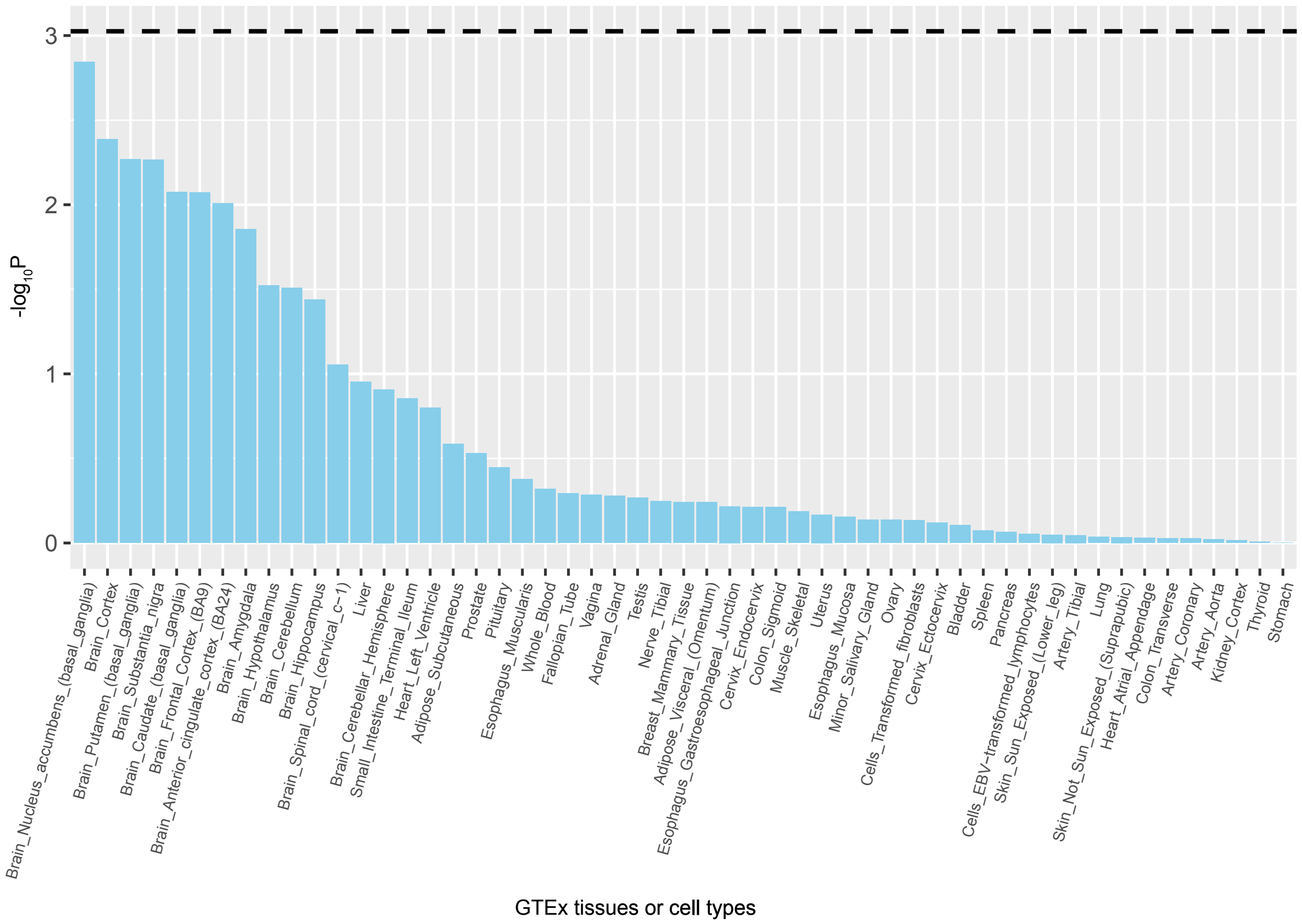

**Supplementary Figure 7. GTEx tissue expression enrichment using MAGMA.** The dashed line is the cutoff for Bonferroni-corrected significance.

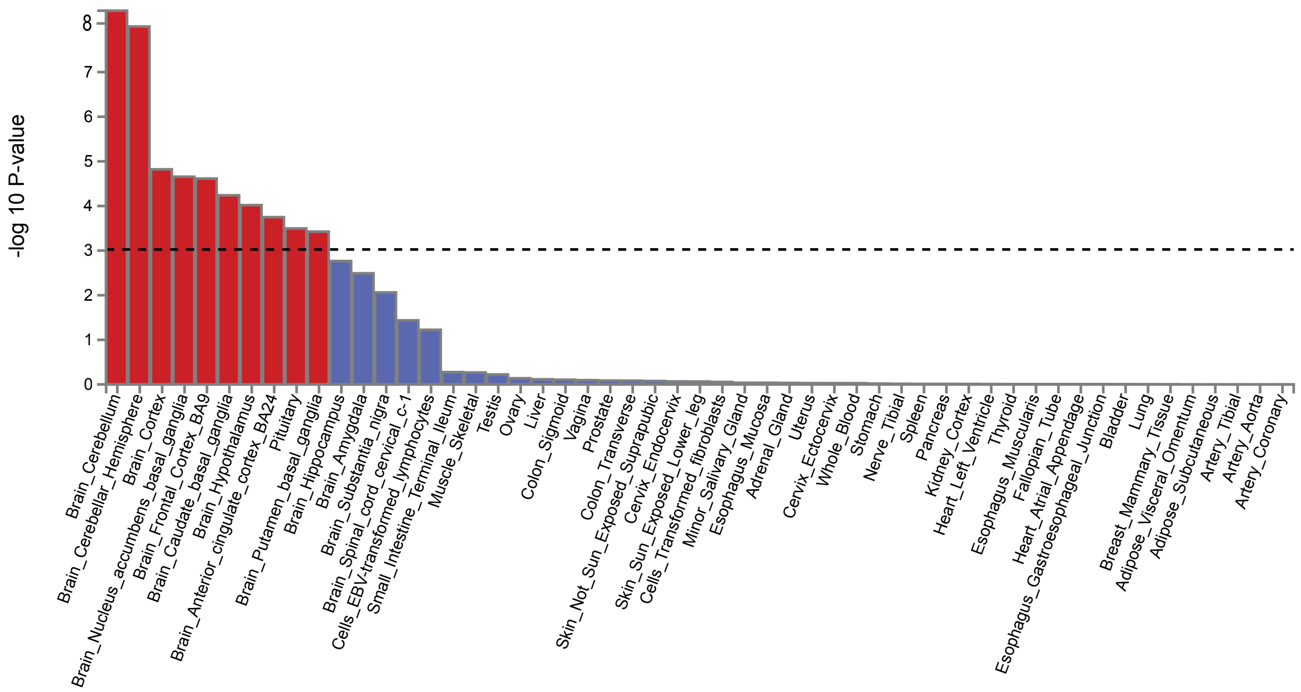

**Supplementary Figure 8. Phenome-Wide associations with PAU PRS in BioVU.**

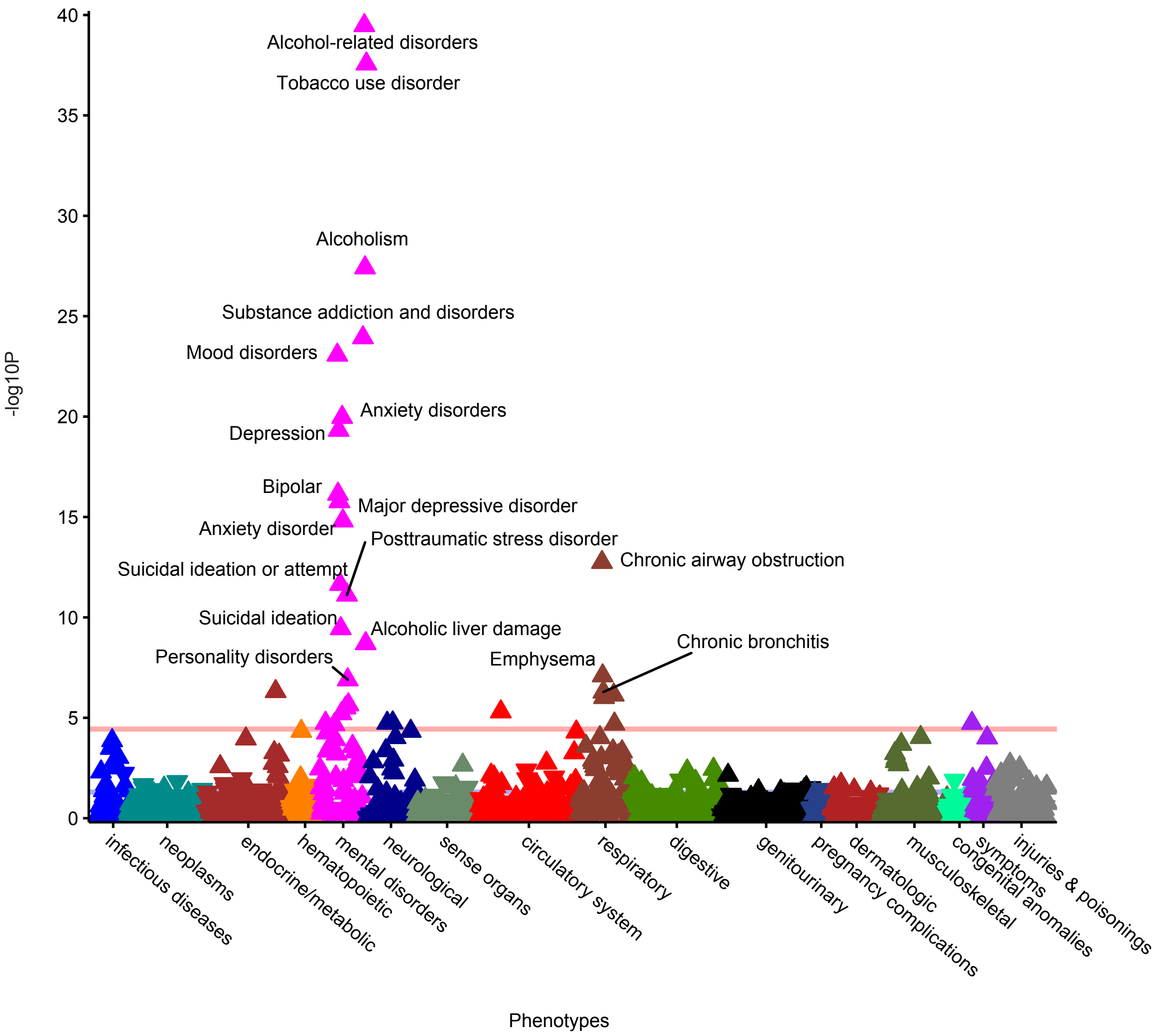

**Supplementary Figure 9. Manhattan and QQ plots for MTAG analysis of PAU.**

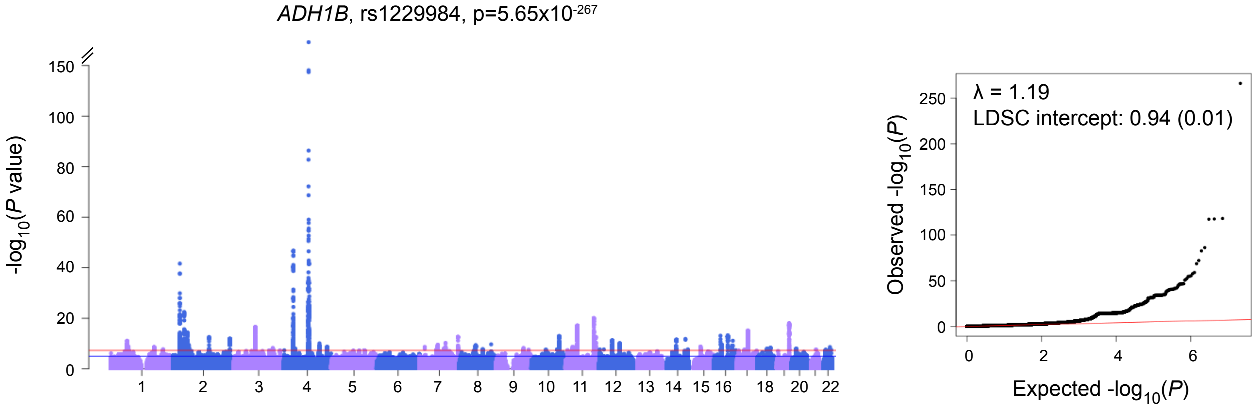

**Supplementary Figure 10. Manhattan and QQ plots for MTAG analysis of DrnkWk.**

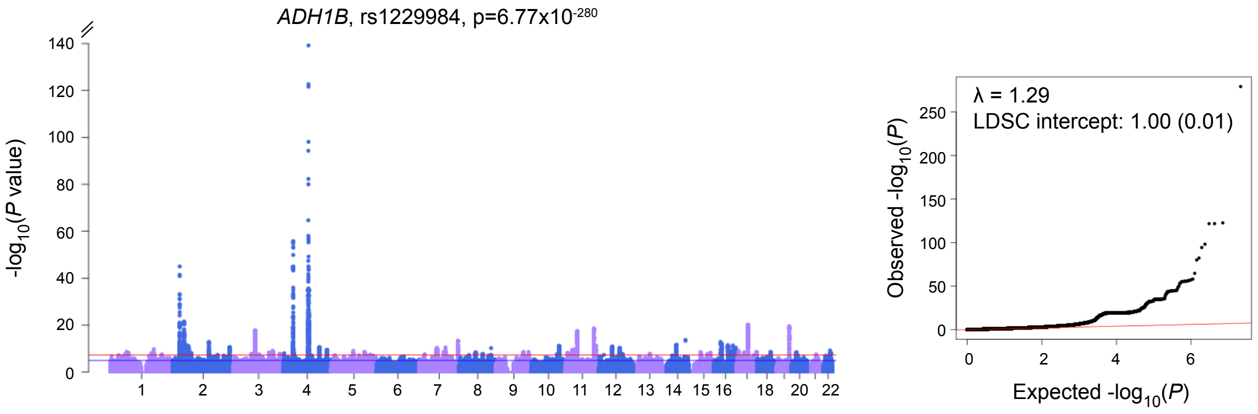
